# Flexible neural encoding predicts the comprehension of degraded speech

**DOI:** 10.64898/2026.06.05.730499

**Authors:** Alexis D. MacIntyre, Tobias Goehring, Matthew H. Davis

## Abstract

How listeners track a variable and continuous acoustic speech signal and parse it into meaningful linguistic representations is a question central to cognitive neuroscience. Moreover, the resilience of this process to acoustic signal degradation is not fully understood. The current study consists of a listening task wherein participants (n = 38) were presented with a naturalistic story whilst undergoing continuous electroencephalography (EEG). Critically, we manipulated access to speech information over two independent dimensions: Spectral clarity, which ranged from unprocessed to severely spectrally degraded; and language, which was either English, spoken by the participants, or Dutch, an incomprehensible language that is closely related to English. All stimuli were produced by the same bilingual speaker. We applied banded regression to model the neural response to a set of acoustic and linguistically derived features. We found that there is no single speech feature for which neural encoding reliably predicted speech understanding in individual listeners or conditions. Yet, differences in feature weights between experimental conditions can be combined into a composite score that strongly predicted individual subjective comprehension of spectrally degraded but linguistically accessible speech in data from held-out participant. Hence, the manner in which neural encoding flexibly adapts to listening context is associated with advantageous perceptual strategies for degraded speech. These findings underscore the complex interplay between low- and high-level features during speech listening and illustrate inter-condition differences in the neural encoding of these properties. Our results highlight natural variation within and between individuals in the encoding of acoustic and linguistic features which provides a potential pathway towards individualised assessment of clinical populations.

## 1 Introduction

Human listeners achieve remarkable success at comprehending speech, despite a wide range of listening challenges. For instance, speech masked by background noise or other voices at adverse signal-to-noise ratios can be readily understood, especially where cues such as pitch are leveraged to segregate target and background sound sources (Brown and Bacon, 2010). Listeners correctly perceive speech content despite dramatic reductions in spectral detail (Faulkner et al., 2000; Davis et al., 2005; Cychosz et al., 2024), harmonic structure (Remez et al., 1981), or temporal structure (Webb and Sohoglu, 2025). Finally, speech understanding is also robust to transient interruptions by silence (Miller and Licklider, 1950), noise (Warren et al., 1997), or other signals (e.g., phoneme restoration; Samuel, 1981; Bashford et al., 1996). Together, these demonstrations of flexibility in speech perception suggest that no single acoustic cue or cues are necessary for comprehension, though many cues may be sufficient for comprehension (Bailey and Summerfield, 1980). Recent work has attempted to index behavioural measures of speech comprehension using neural data; in particular, neural tracking of the speech signal has been proposed as a physiological correlate of understanding (e.g., Iotzov and Parra, 2019). Briefly, neural tracking encompasses a number of related methods to identify the relationship between brain activity and a continuous stimulus over time (Ding and Simon, 2012; Golumbic et al., 2013; Ding and Simon, 2014; Crosse et al., 2016; Wöstmann et al., 2017; Obleser and Kayser, 2019; Molinaro, 2025). One technique is to use temporal response functions to model how speech features (e.g., the amplitude envelope, phonetic transcriptions) map onto time-locked neural responses by estimating a linear filter that best predicts neural measures from the stimulus (forward/encoding model) or vice versa (backward/decoding model; Holdgraf et al., 2017; Kriegeskorte and Douglas, 2019).

Given the resilience of speech perception, however, it is unclear that measures of neural encoding of specific acoustic features can be used to predict individual listening outcomes during connected speech listening (Brodbeck and Simon, 2020; Zoefel and Kösem, 2024). Some studies initially suggested that speech envelope-tracking is modulated by understanding (e.g., Vanthornhout et al., 2018; Etard and Reichenbach, 2019; Chen et al., 2023). Yet, others used manipulations of prior knowledge to modulate perceptual outcomes and have shown that acoustic feature tracking does not necessarily reflect listening outcomes in these situations (Kösem et al., 2023; Karunathilake et al., 2023). It is possible that acoustic feature tracking instead reflects factors that accompany variation in intelligibility, including auditory attention or low-level stimulus properties (e.g., spectral clarity; MacIntyre et al., 2024). Another explanation comes from predictive processing theories in which neural encoding reflects differences between prior knowledge or predictions and sensory signals (Sohoglu and Davis, 2020; Davis and Sohoglu, 2020). In any case, and in line with findings from behavioural studies of speech perception and intelligibility, it does not appear that the neural encoding of any single set of acoustic features can be used to predict comprehension outcomes across listening contexts.

In light of these challenges, measures of the neural processing of higher-level linguistic features have been proposed as a more reliable marker of speech understanding. Unlike acoustic features, which are derived directly from the physical signal, linguistic features (e.g., syllable and word onsets) can be thought of as abstract mental objects with indeterminate acoustic correlates (Twaddell, 1935; Studdert-Kennedy, 1987; Port, 2007). Multiple groups report robust effects of speech comprehension on the neural encoding of linguistic features (Brodbeck et al., 2018; Broderick et al., 2018; Verschueren et al., 2022; Gillis et al., 2023; Tezcan et al., 2023). These studies can involve comparisons of neural responses to intelligible and unintelligible variants of the same stimulus (e.g., normal and sped up speech; Verschueren et al., 2022), a familiar or unfamiliar language (Gillis et al., 2023; Tezcan et al., 2023), attended or unattended speakers in a cocktail party paradigm (Brodbeck et al., 2018; Broderick et al., 2018), and acoustically degraded speech before and after informational priming (Karunathilake et al., 2023). However, these methods may also lead to changes in the degree of auditory attention applied to speech in control conditions. Thus, the extent to which the neural encoding of linguistic markers reflects comprehension success, or rather, attentive listening that includes attempts at comprehension, remains unknown. The nature of the relationship between linguistic encoding and comprehension is also unclear for speech that is incompletely or only effortfully understood. For example, Yasmin et al. (2023) observed that, during noisy speech perception, behavioural measures of intelligibility were uncoupled from semantic feature tracking at low signal to noise ratios.

In this paper we, therefore, explore acoustic and linguistic encoding of clear and degraded speech during story listening across two closely related languages, one of which is understood and the other not understood by experimental participants. To ensure attentive listening throughout, participants are required to detect occasional repeated segments, a task that can be achieved successfully despite signal degradation or lack of language familiarity. This work builds on previous analyses of the same data. Previously, we employed a decoding approach to assess the generalisability of decoders when reconstructing the envelope across differing levels of spectral degradation (MacIntyre et al., 2024). Spectral degradation was applied with a vocoder approach developed and used for the simulation of speech perception with cochlear implants (Grange et al., 2017; Gaultier and Goehring, 2024). We then compared linear to non-linear decoder performance when reconstructing an extended acoustic feature set that also included measures of spectral change over time, finding similar decoding accuracy regardless of model architecture (MacIntyre et al., 2026). This latter study also incorporated extensive random permutation testing, which showed that certain acoustic features are more robustly decoded than others. However, decoder-based analyses are limited, in that it is not possible to combine multiple neural predictors within one model. This precludes feature importance ranking. Moreover, decoders are less well suited to linguistic features, typically represented as binary vectors, on account of their sparseness (Gillis et al., 2022).

Here, we use expectation maximisation (Fuglsang et al., 2024) to fit complex encoding models that combine both acoustic and linguistic predictors of neural activity. We perform factorial analyses of condition-specific model weights relative to null distributions to evaluate how neural encoding of acoustic and linguistic features change with speech clarity for comprehended and non-comprehended languages. These analyses indicate that there is no single acoustic or linguistic feature that is necessarily encoded during comprehension or that is sufficient to distinguish comprehended and non-comprehended speech at all levels of clarity. Since neural encoding of single acoustic or linguistic features cannot be used to predict comprehension, we developed a data-driven method that combined multiple measures of neural encoding to predict subjective ratings of listening outcomes for held-out participants. This analysis used repeated resampling of 50% splits of the data set to ensure that neural predictors of comprehension performance generalise across subjects. Thus, our approach captures both a stable underlying component capable of describing group-level behaviour, as well as a path towards listener profiling by identifying where individuals are positioned along this axis (Halai et al., 2017). Our results show that by contrasting neural encoding of acoustic and linguistic features in different listening conditions we can accurately predict listening outcomes in unseen participants. This finding indicates that the flexibility of neural encoding of acoustic and linguistic features both predicts resilient speech comprehension in the face of challenging listening situations, and provides cross-subject, generalisable neural markers of that resilience. These findings suggest that comparisons of different listening conditions and weighted combinations of acoustic and linguistic feature encoding are required to predict individual speech comprehension from neural measures.

## 2 Methods

Complete details concerning the participants and stimuli generation and processing are available in MacIntyre et al. (2024) and MacIntyre et al. (2026). Briefly, the data set consists of thirty-eight typically hearing adults (22 female and 16 male; aged 18–35, Mean 24.95, SD 5.09). Informed consent was obtained from all participants for being included in the study. The study received approval from the Cambridge Psychology Research Ethics Committee and was conducted in accordance with the Declaration of Helsinki. The speech stimuli consisted of the original English and Dutch translation of the Sherlock Holmes story “The Adventure of Charles Augustus Milverton” (Doyle et al., 1903; Doyle, 1903), performed by a bilingual adult male who grew up speaking Dutch at home and American English at school. The English story was 33 min. 47 s and the Dutch story was 35 min. 54 s, each split into six trials with two trials per experimental condition. We created the vocoded conditions using SPIRAL vocoder (Grange et al., 2017) with two levels of spectral degradation: The first contained 16 frequency analysis bands or channels, resulting in moderately spectrally degraded but highly intelligible speech. We refer to this condition as “vocoded”. The second level was also formed from 16 analysis bands, but the edges of the bands overlap by −16 dB per octave to create the “vocoded + blurring” condition, which is more severely spectrally degraded.

### 2.1 Task and Behavioural Measures

**Repeated Phrase Auditory Detection Task** It is established that attended speech evokes a neural response distinct from that of non-attended speech (e.g., Ding and Simon, 2012; Rimmele et al., 2015; Biesmans et al., 2016; Vanthornhout et al., 2019; Geirnaert et al., 2021; Belo et al., 2021). This is potentially problematic where an effect of interest, such as speech intelligibility, is likely to co-vary with attention. Hence, to promote attentive listening across the experiment, we introduced an auditory detection task. Participants were instructed to listen for short sections (Mean 2.06 s, SD 0.04) of the speech audio that repeated once before continuing with the story, and to press a button upon detection. The repeated audio segments always encompassed a complete sentence or clause demarcated by short natural pauses, which enabled us to splice the recording without producing acoustic artifacts or other possible cues. We recorded hits (correctly identified targets), false alarms (incorrectly identified targets), and response times measured from the onset of the repeated phrase.

**Behavioural Ratings** After each trial, participants were prompted to rate the preceding speech on a seven-point Likert scale according to how well they could follow the story and how engaged they felt by the story content, where 1 = no understanding or engagement and 7 = full understanding and engagement. We clarified with text and verbal instructions to participants that ability to follow, in this case, meant comprehension of the plot and dialogue. Engagement was defined as feeling emotionally invested in learning the outcome of the story. Since engagement largely tracked with understanding, we do not analyse engagement further.

**Procedure, EEG Acquisition, and Preprocessing** The experiment was run in Psychtoolbox (Kleiner et al., 2007). We recorded 64-channel EEG at 2048 Hz using a BioSemi ActiveTwo EEG system (BioSemi, Amsterdam, The Netherlands) with scalp electrodes positioned following the International 10-20 scheme. Participants heard the stimuli binaurally from ER-2 insert earphones (Etymotic Research, Elk Grove Village, USA). We performed preprocessing of the EEG in MATLAB (MATLAB, 2020) using the Fieldtrip (Oostenveld et al., 2011) and noisetools (de Cheveigné and Arzounian, 2018) toolboxes. Data were re-referenced to the average and downsampled to 256 Hz. After removing residual line noise (Klug and Kloosterman, 2022), we applied a highpass filter at 0.5 Hz and then a lowpass filter at 40 Hz with 4th order Butterworth filters. Noisy or flatlining channels were visually identified and subsequently interpolated based on neighbouring electrodes. We removed eye blink artifacts using independent component analysis (runica algorithm). The first and final 5 seconds of each trial were excluded to omit potential filtering artifacts Crosse et al. (2021). Finally, we down-sampled the EEG and features to 100 Hz for the encoding analysis. For the purposes of model selection and statistical comparisons across experimental conditions, we identified predetermined channel groups covering regions of interest identified in previous speech perception studies (e.g., Montoya-Martínez et al., 2021; Simon et al., 2022). We refer to these as central frontal (CF; Fp1, AF3, Fpz, Fp2, AF4, Afz), left frontal (LF; F3, F5, F7, FT7, FC5, FC3), right frontal (RF; F4, F6, F8, FT8, FC6, FC4), central posterior (CP; P1, P03, POz, Pz, P2, PO4), left posterior (LP; P3, P5, P7, P9, PO7, PO3), and right posterior (RP; P4, P6, P8, P10, PO8, PO4).

### 2.2 Analysis

**Acoustic Feature Extraction and Processing** The speech acoustic feature set can be sub-categorised as primarily pertaining to either the amplitude envelope or the fine structure (spectral properties) of the signal. Starting with the amplitude envelope itself, we generated this feature by passing the unprocessed and vocoded stimuli audio through an Equivalent Rectangular Bandwidth (ERB) filterbank comprising 32 filter analysis bands spanning 50 to 5000 Hz. The filtered output was rectified, power law-compressed, and averaged across bands (Biesmans et al., 2016). The other envelope-based feature was the first derivative of the positive-going envelope, which we term “Envelope Derivative“. This acoustic landmark is sometimes also referred to as “acoustic edges” (Doelling et al., 2014). It roughly coincides with phonetically defined vowel onsets in speech (MacIntyre et al., 2022) and has been proposed as an important cue during both speech and music perception (Ding and Simon, 2012; Doelling et al., 2014; Oganian and Chang, 2019; Leske et al., 2025).

The spectral features were generated using the Timbre Toolbox (Peeters et al., 2011). Although this software package is most commonly used within psychophysics and music information retrieval, many of the spectral descriptors provided therein have also been applied across speech and voice research (e.g., Lauraitis et al., 2020; Taylor et al., 2020; Sandhya et al., 2020; Pommée et al., 2021; Schultz and Vogel, 2022). We extracted the features from the output of an ERB filter bank with the same parameters used to generate the envelope. The pipeline was configured to operate on a window of length 35 ms with hop of 10 ms. To improve stability and reduce noise at lower signal levels, we scaled the resulting features by a signal intensity threshold with Gaussian smoothing. For efficiency, we pre-emptively removed some features that were highly correlated with others: These were spectral kurtosis (spectral skew *r* 0.95), spectral flatness (spectral spread *r* 0.85), and spectral roll-off (spectral centroid *r* 0.87). The acoustic features retained for model selection are summarised in Table 1 and visualised in Figure 1, Panel C.

**Figure 1:**
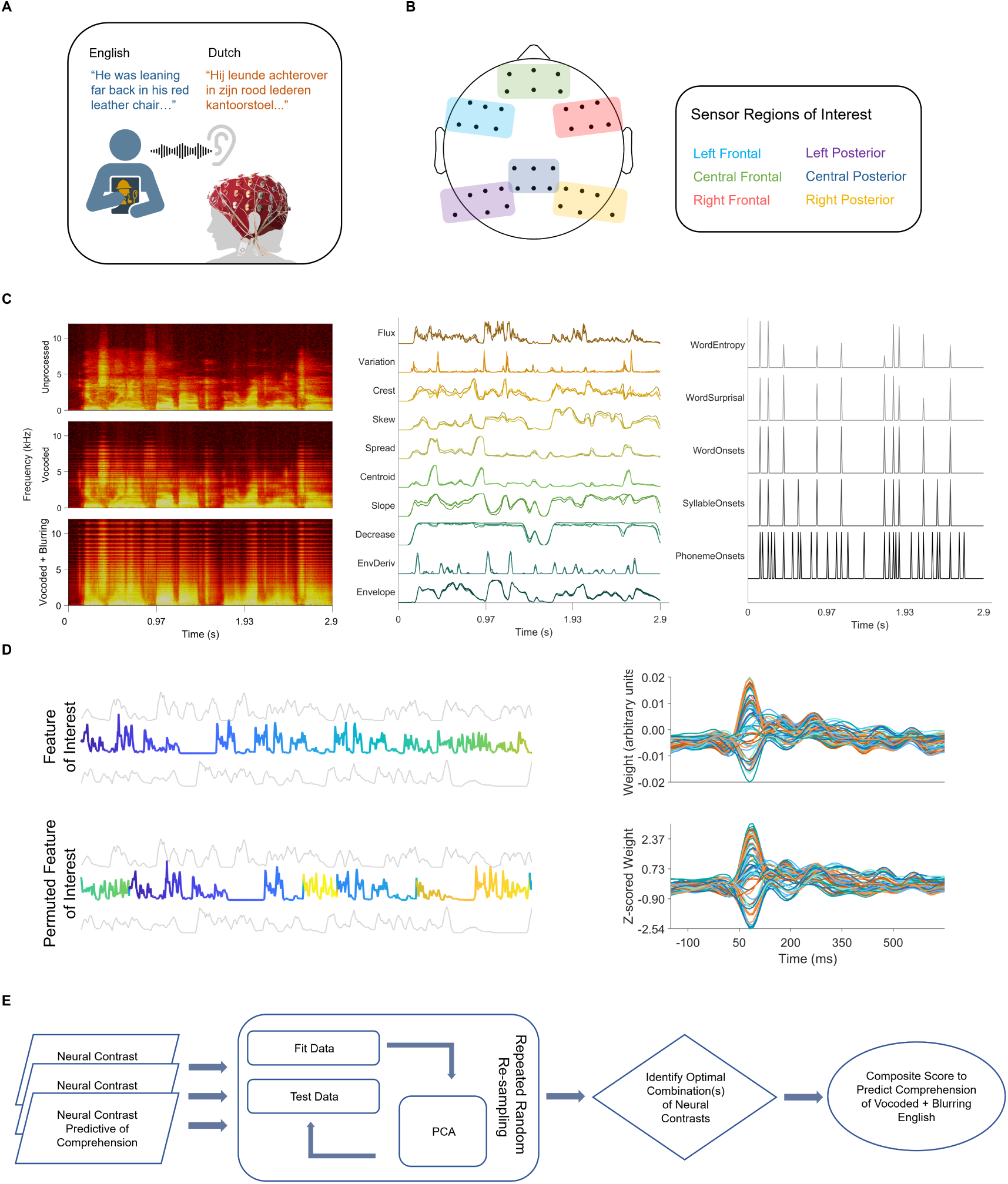
Panel A: Exemplar text materials in English and Dutch with experimental set-up. Panel B: Schematic diagram depicting the electrode or sensor regions of interest. Panel C: Spectrograms (left panel) showing a speech segment at each level of spectral degradation. Acoustic features (middle panel) with spectral degradation conditions overlaid. Linguistic features (right panel) with spectral degradation conditions overlaid. Panel D: Illustration of the procedure for generating *Z*-scored weights. The left panels depict a highlighted feature of interest (top) and its randomly permuted form (bottom), with faded grey traces indicating other features, which are held constant. The right panels show the associated weights for that feature in their original (top) and *Z*-scored (bottom) forms. Each individual trace corresponds to an EEG sensor. Panel E: Flowchart summarising the exploratory individual differences principal component analysis (PCA) to predict comprehension of vocoded + blurring English.

**Table 1:** Features from The Timbre Toolbox (Peeters et al., 2011) applied within the neural encoding model selection.

| Feature | Description | Related Terms and Concepts |
| --- | --- | --- |
| Amplitude Envelope | Instantaneous amplitude of the signal |  |
| First Derivative of the Envelope | Peaks in the rate of change of the amplitude envelope | Acoustic edges, attack, rise time, peakRate |
| Spectral Centroid | Amplitude-weighted spectral mean | Spectral centre of gravity, brightness, first spectral moment |
| Spectral Spread | Dispersion of the spectrum around the centroid | Second spectral moment |
| Spectral Skew | Asymmetry of the spectrum around the centroid | Spectral tilt, third spectral moment |
| Spectral Crest | Ratio between the mean and maximum of the spectral magnitude |  |
| Spectral Slope | Linear fit of the magnitude spectrum across frequencies |  |
| Spectral Decrease | Conveys the reduction of spectral magnitude with increasing frequency |  |
| Spectral Variation | Dissimilarity of the frequency spectrum over time |  |
| Spectral Flux | Absolute difference in the magnitudes of consecutive spectra | Cochlea-scaled entropy |

**Linguistic Feature Extraction** We aligned the text transcriptions to the speech recordings using the WebMAUS forced aligner (Kisler et al., 2012) and extracted word, syllable, and phoneme onsets, which we manually inspected and corrected where necessary. Word surprisal is a measure of how unexpected a word is in a given context, defined as the negative logarithm of its conditional probability given the preceding words. Similarly, word entropy quantifies the overall uncertainty about which word will appear next, and is defined as the expected surprisal across all possible next words (Levy, 2008; Gwilliams and Davis, 2022). We estimated surprisal and entropy using large language models (GPT-2; Radford et al., 2019; Wolf et al., 2020; Boeve and Bogaerts, 2025), incorporating the preceding 30 tokens (Tezcan et al., 2023). The Dutch transcript was processed by a GPT-2 model re-tuned for the Dutch language (de Vries and Nissim, 2020). The time-stamps for the linguistic onsets were converted into one-hot vectors, and the Word surprisal and entropy predictors were used to scale the word onset vector. An example of the linguistic features is shown in Figure 1, Panel C.

**Encoding models** We investigated the neural response to speech features using encoding models individually fit within participant and experimental condition, covering-150 to 650 ms relative to stimulus onset. When estimating a linear encoder, some form of regularisation is often used to shrink the coefficients of uninformative or noisy predictors (e.g., by using a ridge estimator; Crosse et al., 2016). However, the resulting weights may still be unreliable when features are linearly dependent on one another, which is typical of naturalistic stimuli such as speech (Nunez-Elizalde et al., 2019; La Tour et al., 2022). Here, we employ expectation maximisation-or EM-banded regression to estimate the TRF as introduced by Fuglsang et al. (2024). This approach incorporates an algorithm to shrink or regularise “bands” or groups of predictors by iteratively refining parameter estimates based on the data. Briefly, each feature band shares a common amount of regularisation, which controls how strongly the weights are pushed toward zero. The EM algorithm alternates between estimating the likely values of the regression weights, then updating the group-level shrinkage strength to better fit the observed data. This cycle continues until stable values are reached (Do and Batzoglou, 2008). The resultant encoding weights, sometimes called the temporal response function (TRF), are interpretable as an index of neural activity over [sensor-]space and time, analogous to a classical event-related potential (Crosse et al., 2021). Simulations suggest that, in comparison with ridge regression, EM-banded regression is effective at recovering the appropriate weights for informative features whilst shrinking those of potentially correlated, yet uninformative features (see Fuglsang et al., 2024, for further discussion and caveats). We optimised the smoothness parameter *H* for each predictive feature individually using 4-fold cross validation where *H* could range from 0 to 24 with a step size of 4. For acoustic features, *H* was 0 (Envelope and Spectral Flux) or 4 (Envelope Derivative and Spectral Variation), and for linguistic features, *H* was 16. We left all other tunable parameters as the default values (Fuglsang et al., 2024).

**Acoustic feature selection** We expect that different acoustic features, including spectral descriptors, will correlate with the envelope (Shu et al., 2016; Oxenham et al., 2017). Therefore, to encourage model parsimony whilst optimising predictive power, we performed stepwise model comparisons to determine which acoustic features to include in the model (Kriegeskorte and Douglas, 2019). Evaluations were performed on 60 seconds of held-out test data from the middle of the experiment (Crosse et al., 2021). Starting with single-predictor models, we determined a baseline model based on the feature that achieved lowest mean-normalised root mean square error (nRMSE). We then added the remaining acoustic features one at a time, taking forward the model that improved *r* with the lowest nRMSE. Both *r* and nRMSE are averaged across experimental conditions and regions of interest. Note that, when calculated using out-of-sample data, *r* and nRMSE can rank models differently (Murphy, 1988). We consider both metrics here as nRMSE reflects estimator bias and variance, and *r* provides a sense of how well the encoding model’s predictions track patterns in the observed neural data over time (irrespective of magnitude errors). We compared candidate to baseline model *r* and nRMSE using two-tailed signed rank tests. When we ceased to find improvements in *r* without deleteriously affecting nRMSE, the acoustic part of the encoding model was finalised. A summary of the procedure is shown in Figure 1, Panel D. We subsequently included the linguistic onsets as a set, and then word entropy and surprisal. As we had an *a priori* interest in comparing acoustic to linguistic features, we did not base our decision on significance testing for improved *r* in this case, although we do report the test results.

**Random permutation of model weights** For robustness and to allay nuisance differences across predictors and participants, we scaled the encoding weights relative to an empirically determined null distribution (i.e., a distribution of weights where no true stimulus-brain relationship exists). We executed this step by holding all other features in the model constant and randomly shuffling ∼ 2 s segments of the feature of interest, before concatenating them to form a randomised surrogate (Figure 1, Panel E). This method preserves the local temporal structure of the feature, ensuring its plausibility, but destroys long-term correspondence to the neural response. Segments were split at signal zero-crossing points and joined using smooth interpolation to ensure that the shuffled feature was free of artifacts. Within-participant, and for each feature and listening condition, we performed 250 iterations on a high-performance computing cluster.

#### 2.2.1 Planned group-level analysis

**Statistical testing** To model the mean response within each region of interest, we ran linear mixed effect models (MATLAB function fitlme fit with restricted maximum likelihood) containing fixed effects of *Language* × *Spectral Degradation* with random intercepts for *Participant* and *Electrode* over each lag between −100 and 600 ms. The two levels of Language are English and Dutch, and the three levels of Spectral Degradation are unprocessed, vocoded, and vocoded + blurring. The model reference level was unprocessed English.

To identify statistically significant differences over time, we first applied cluster-based permutation tests with 5000 iterations (Maris and Oostenveld, 2007). Temporal clusters within sensor regions were defined for consecutive 10 ms time windows with *t*-statistics ≥ 1.96, and were considered significant after adjusting the permutation-based alpha for repeated comparisons (i.e., Bonferroni-corrected for six channel groups; *p* ≤ 0.05*/*6 = 0.008). The minimum length of a cluster was arbitrarily set at a threshold of 50 ms, and the cluster maximum statistic must also reach *t* ≥ 3.291 (approximating a two-tailed test at *p* = 0.001). We confirmed the presence of specific experimental effects using F-tests with Satterthwaite degrees of freedom (MATLAB function coefTest)

After identifying significant temporal clusters that met our threshold criteria, we confirmed each linear mixed effect model associated with the cluster maximum and obtained pairwise contrasts across experimental conditions in RStudio (Posit team, 2025) using functions from the lme4 (Bates et al., 2015), car (Fox and Weisberg, 2019), and emmeans

(Lenth and Piaskowski, 2025) packages. For the post-hoc tests, alpha was determined according to nine planned pairwise contrasts: English versus Dutch within the three levels of spectral degradation, and all pairwise comparisons between levels of spectral degradation within the two languages (*p* ≤ 0.05*/*9 = 0.006). The 95% confidence intervals provided for descriptive statistics are generated using the bootstrap method with resampling and 5000 iterations.

#### 2.2.2 Data-driven analysis of individual differences in comprehension

In conducting the planned analysis, we incidentally observed that neural encoding in certain conditions covaried with encoding in other conditions. This led us to hypothesise that within-participant differences across conditions could be more informative than the raw encoding weights in isolation. Shared variance across feature encoding provides the opportunity to reduce dimensional space. Hence, by combining multiple measures, we can potentially improve the signal-to-noise ratio in the analysis, as well as identify an underlying latent component that more succinctly summarises an outcome of interest in the data set. Principal component analysis (PCA; Jolliffe, 2004) is well suited to this purpose: PCA can identify latent dimensions that capture shared variance among predictors and is commonly used for feature reduction and to generate composite scores or indices (Mwangi et al., 2014). Here, we examined whether a composite score combining multiple neural contrasts would form a more powerful predictor of comprehension in comparison to its constituent parts (Figure 1, Panel E).

We target understanding of vocoded + blurring English, as self-reported comprehension of unprocessed and vocoded English was around ceiling level for most participants. To baseline-correct for differences in Likert scale ratings, we calculated the understanding measure (henceforth,’comprehension’) by subtracting comprehension ratings for vocoded + blurring English from those for unprocessed English. We then generated neural contrasts, for each feature and sensor region of interest, across all pairwise comparisons that varied on a single dimension (e.g., Vocoded + blurring English − vocoded + blurring Dutch; see Supplementary Material, Table 11 for complete list). These contrasts were calculated by dividing the encoding time course (−100 − 600 ms post-stimulus onset) into overlapping 40 ms windows with 10 ms step size. Depending on whether the reference condition had positive or negative weights on average, we subtracted the maximum of the comparison condition (e.g., unprocessed Dutch) from the maximum of the reference condition (e.g., unprocessed English), or subtracted the minimum of the comparison condition from the minimum of the reference condition. The resultant contrasts were then correlated to comprehension using Spearman’s rho. As an initial feature reduction step, we performed cluster-based permutation tests over time windows to exclude transient or spurious correlations. We first set an arbitrary threshold where each cluster had to persist across at least three adjacent windows where rho ≥ 0.30, and achieve a cluster maximum rho ≥ 0.40. We then randomly permuted the correlations over time with 1000 iterations and accepted the observed cluster if *p_cluster_* ≤ 0.008 (based on multiple comparisons over six regions of interest). Where the same contrast and speech feature re-appeared across multiple regions of interest at roughly the same time points, we retained only the region with the highest cluster sum rho. For each cluster, the time window where the maximum rho occurred was taken forward as a neural contrast.

To determine whether the neural contrasts could then be mapped onto a common underlying component, we applied PCA. We confirmed stability of the loadings and that the resulting principal components were robust by performing all analyses using repeated split-half resampling. Specifically, we randomly divided the complete data set in two (*n* = 19 participants per group), fitting the PCA to just one half. The held out data in the other half were normalised using the mean and standard deviation of the fit data, and projected into the fit principal component space. We then correlated the first principal component of the test data set with comprehension. The split-halves were randomly re-allocated and the process repeated to produce a distribution of held-out correlations with comprehension (*n* = 10, 000 iterations). From an initial set of 30 neural contrasts that survived cluster-based permutation testing, we evaluated groups of differences ranging in size from four to fifteen neural contrasts (the upper limit of computational feasibility). To identify which inputs to include in the final PCA, we took the highest median rho across these combinations of neural contrasts. This finalised solution produces an emergent first principal component, which we then used to correlate with comprehension of vocoded + blurring English speech. We report descriptive statistics characterising the distribution of held-out rho (i.e., fit and tested on independent split-halves) as well as rho when the PCA is fit to the complete data set.

To summarise, we: (1) Generated contrasts of neural encoding weights between pairs of conditions; (2) Used these contrasts to obtain correlations predicting comprehension of degraded speech during familiar language listening; (3) Combined data from multiple distinct time-periods, sensor regions, and encoding contrasts that each correlated to self-reported understanding using PCA; and (4) Tested whether a principal component derived from these neural measures predicted comprehension ratings.

## 3 Results

Behavioural ratings and results of the repeated phrase auditory detection task are reported in full in MacIntyre et al. (2024), with comprehension ratings and response times to auditory targets visualised in Figure 2. Briefly, on a scale from 1 − 7 where 1 = no comprehension and 7 = perfect comprehension, ratings for English conditions ranged from Median 6.50 (IQR = 1.00) for Unprocessed, to 5.00 (IQR = 2.00) for Vocoded + Blurring. The equivalent values for Dutch were Median 1.25 (IQR = 1.00) for Unprocessed, and 1.00 (IQR = 0.88) for Vocoded + Blurring. The fastest responses were to unprocessed English targets (Mean = 1.03 s, SD = 0.33) and slowest to Dutch Vocoded + Blurring (Mean = 1.39 s, SD = 0.45). Poorer comprehension was predictive of slower response times in the vocoded + blurring English condition (Rho = −0.52, *p* = 0.001, 95% CI [−0.56, −0.16]), thereby providing a corroborating objective measure to the self-reported data (Figure 2, Panel B).

**Figure 2:**
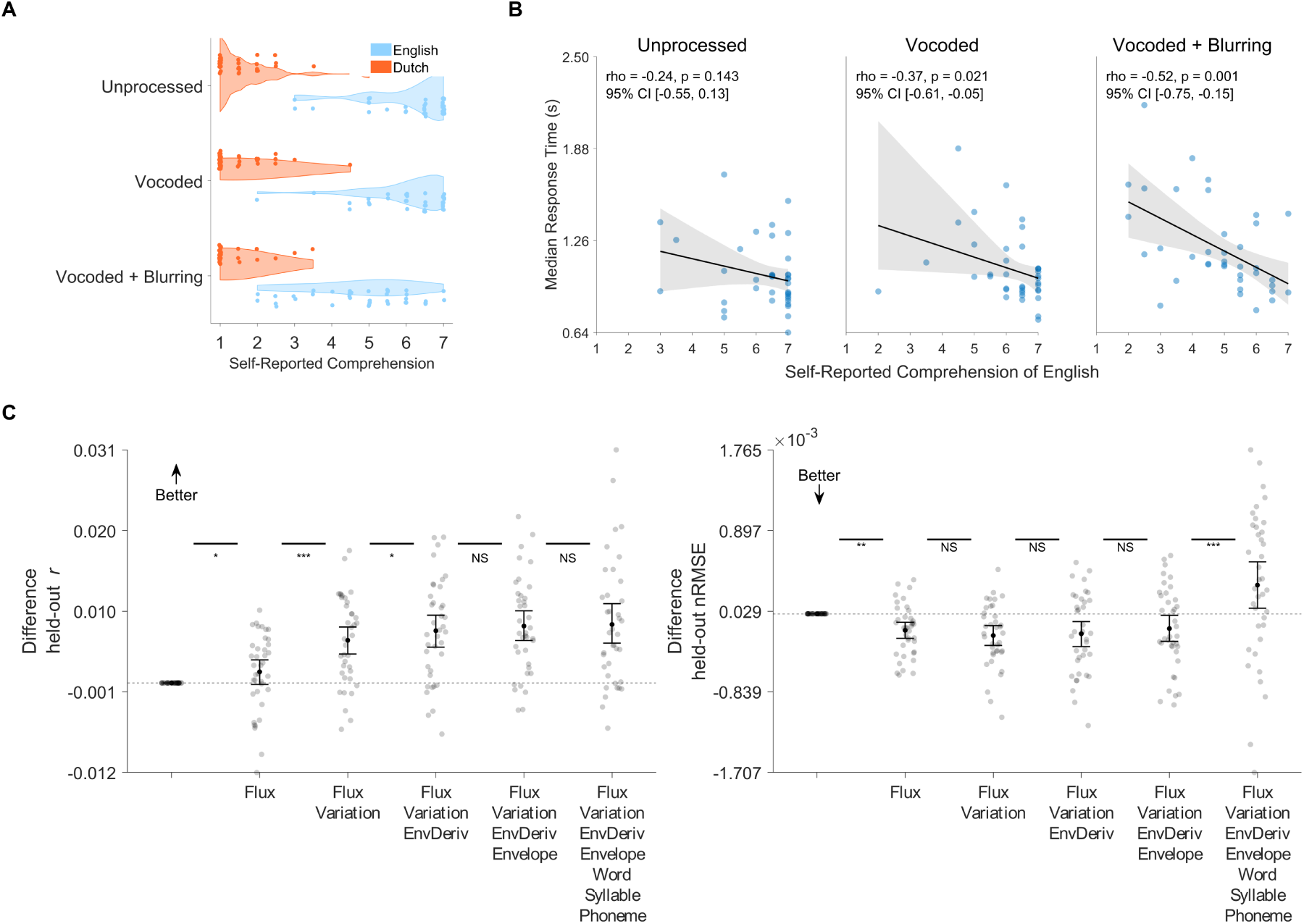
Panel A: Self-reported ratings of speech comprehension across experimental conditions, where 1 = no understanding and 7 = perfect understanding. Ratings are averaged across two blocks per condition. Panel B: Median response times to auditory targets in the repeated phrase detection task as a function of self-reported comprehension of English experimental conditions. Lines and shaded regions indicate the least-squares linear fit with bootstrap 95% confidence intervals. Panel C: Summary of difference in out-of-sample *r* (left panel) and normalised-root mean square error (right panel) in the stepwise model selection procedure. The data are plotted as a difference from the baseline Envelope-only model. Significance bars indicate the results of signed rank tests comparing between nested models, NS *not significant*, * *p ≤* 0.05, ** *p ≤* 0.01, *** *p ≤* 0.001.

### 3.1 The Listening Brain Tracks Spectral Change Alongside the Amplitude Envelope

We adopted a stepwise selection procedure to assess which acoustic features to include in the encoding model (Supplementary Material, Figure 1). Our model selection criteria were improvement in out-of-sample *r* without increasing out-of-sample nRMSE (see Methods for details). The results are summarised in Figure 2, Panel C. The best single-predictor encoding model was Spectral Flux, which conveys differences in the spectrum over time and is sensitive to both frequency and amplitude. Spectral Flux was recently implicated in the neural processing of both speech (Giroud et al., 2024) and music (Weineck et al., 2022) and is conceptually similar to cochlea-scaled entropy (Stilp and Kluender, 2010; Aubanel et al., 2018). Compared to an encoding model containing only Envelope, Spectral Flux has significantly higher *r* (*z* = 2.11, *p* = 0.035) and lower nRMSE (*z* = −3.08, *p* = 0.002).

To the model containing Spectral Flux, the best additional predictor was Spectral Variation. Like Spectral Flux, Spectral variation is a measure of spectral change over time. However, it is invariant to changes in loudness and can be considered more distinct from Envelope than Spectral Flux, which co-varies with signal intensity. Spectral Variation significantly improved *r* (*z* = 3.73, *p* = 0.002) and did not increase nRMSE (*p* = 0.871). The next feature to be included in the Spectral Flux + Spectral Variation model was Envelope Derivative, which improved *r* (*z* = 2.43, *p* = 0.015) without negatively impacting nRMSE (*p* = 0.627). This result supports previous work implicating that acoustic landmark in the neural tracking of speech (Ding and Simon, 2014; Oganian and Chang, 2019). Finally, we found Envelope numerically improved *r*, though this was non-significant (*z* = 1.46, *p* = 0.144). Envelope did not increase nRMSE (*p* = 0.144), but the remaining acoustic features did, either significantly or as a trend (range *p <* 0.001 to 0.074). Given the conventional use of the Envelope throughout the literature, we opted to include it in the model, but ceased evaluating further acoustic features thereafter. Note that, using regularised ridge regression, we found the same four acoustic features produced the highest predictive decoding accuracy in our previous analysis (MacIntyre et al., 2026).

We next introduced the linguistic onset features, which improved held out *r* numerically, though not statistically (*p* = 1.00), but did increase nRMSE (*z* = 3.73, *p* = 0.002). In light of our interest in comparing linguistic to acoustic speech properties, we opted to include this feature set in the final model. However, the additions of Word Surprisal and Word Entropy actually worsened *r*, in addition to deleteriously affecting nRMSE. In case this effect was due to differences in comprehension, we inspected the unprocessed English condition only and found the same result as when collapsing across conditions. Given the additional computational cost of including them, we did not proceed with this most abstract feature set. In sum, the final encoding model contained Spectral Flux, Spectral Variation, Envelope Derivative, Envelope, and Phoneme, Syllable, and Word Onsets. We plot feature weights for the unprocessed English condition in Figure 3 by sensor region of interest. The other experimental conditions are shown in Supplementary Material, Figures 2-6.

**Figure 3:**
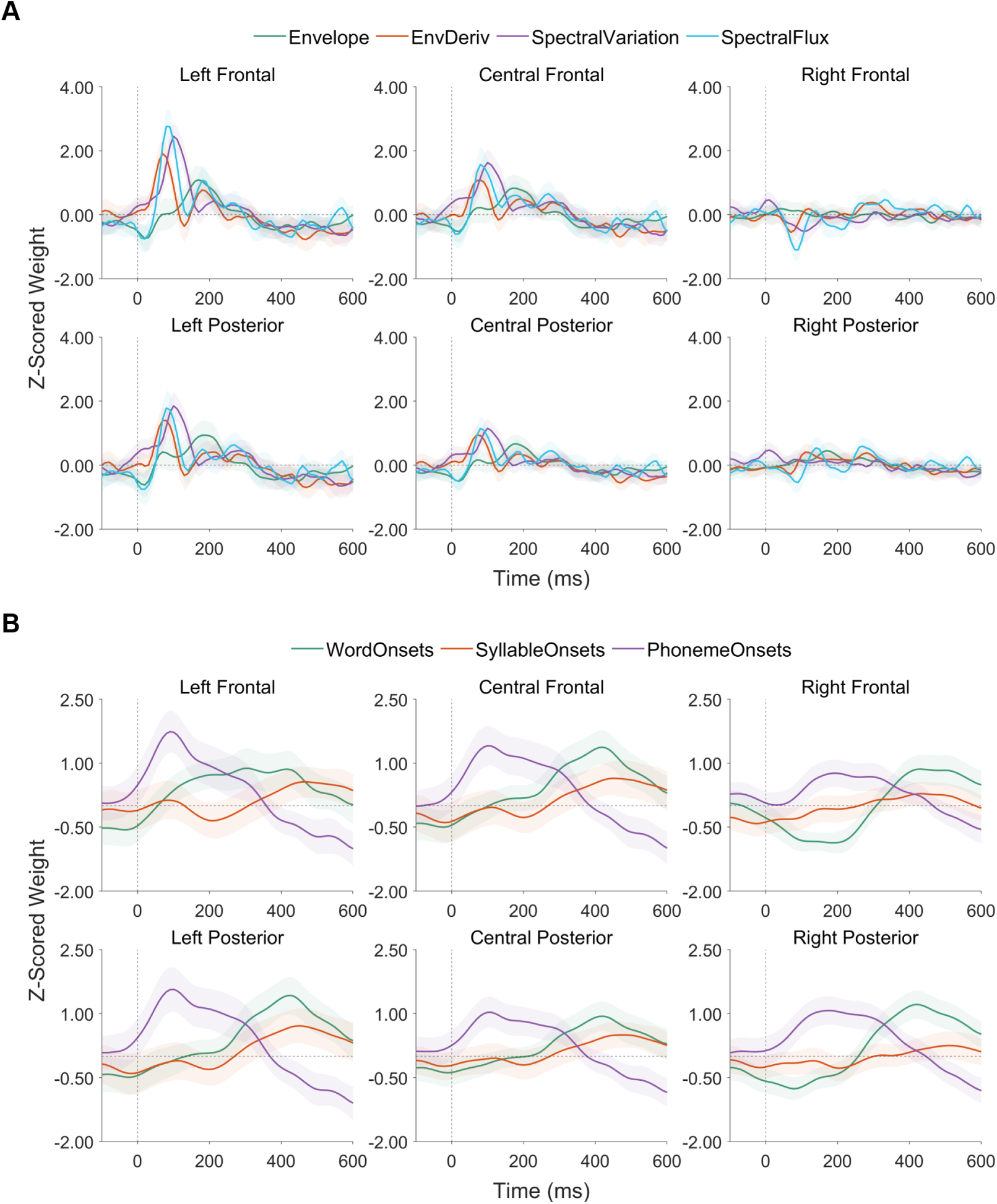
Z-scored encoding model weights for unprocessed English by sensor region of interest. Traces and shaded regions indicate the group mean (averaged over sensors) and 95% confidence intervals of the mean. Panel A: Acoustic features; Panel B: Linguistic features.

### 3.2 Effects of Language and Spectral Degradation on Neural Encoding

We performed linear mixed effects modelling to assess whether acoustic and linguistic feature encoding for comprehended and non-comprehended languages across differing levels of clarity. In the interest of brevity, we will focus on a subset of representative effects (Figure 4) that are sustained and recur over multiple sensor regions. Details concerning selected linear mixed effect models are available in Supplementary Material, Section 6. A complete list of all significant clusters and plots for every combination of ROI and feature are also reported in Supplementary Material, Table 1 and Figures 7–13.

**Figure 4:**
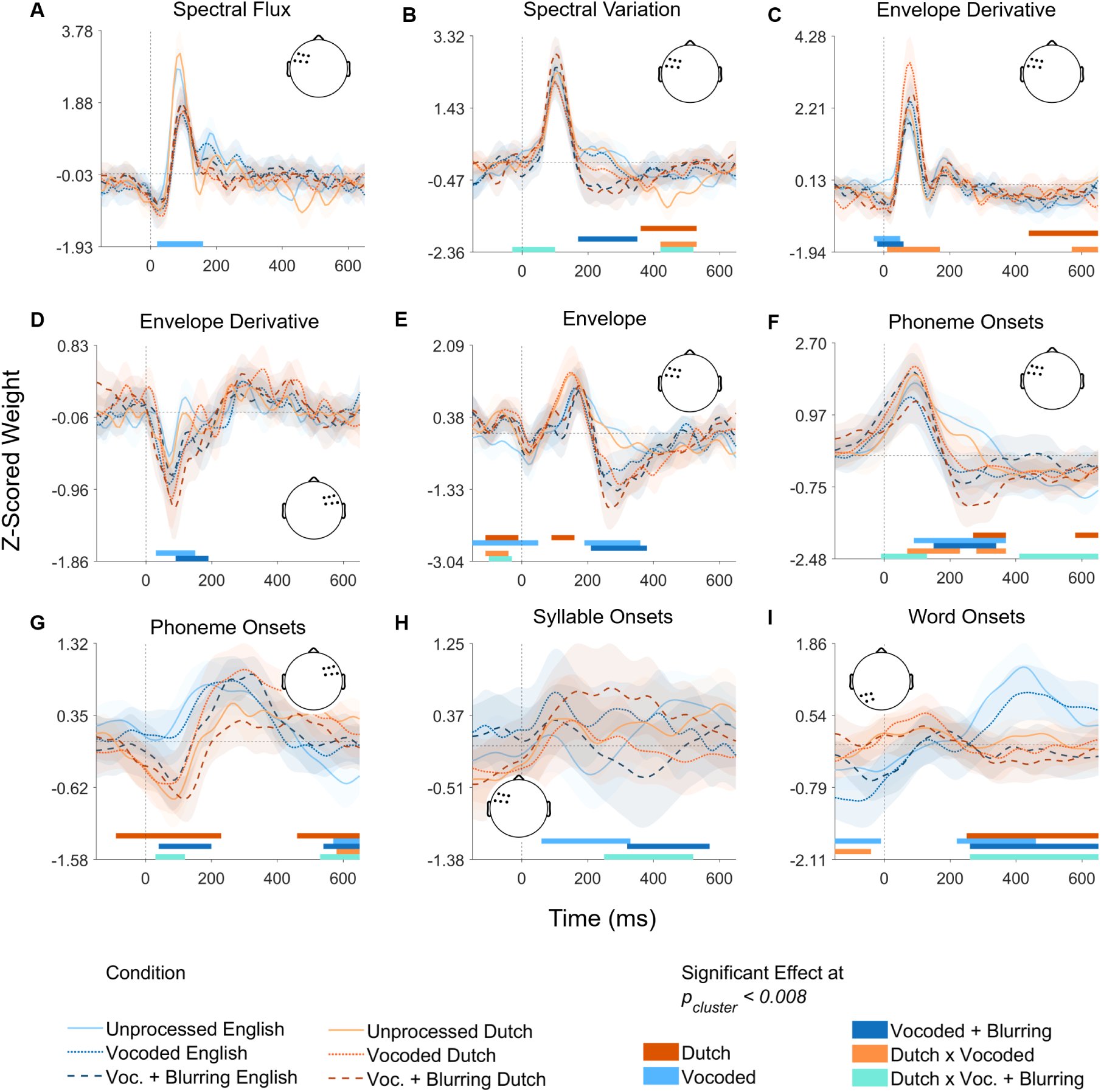
Selected Z-scored encoding model weights, averaged within sensor region of interest, across experimental conditions for speech features with cluster-level results for linear mixed effect models. The model reference level is unprocessed English. Coloured bars indicate time points where a significant effect was found according to cluster-based permutation testing (5,000 iterations). Head diagrams indicate the sensor region of interest. Traces and shaded regions indicate the group mean (averaged over sensors) and 95% confidence intervals of the mean.

#### 3.2.1 Acoustic Features

Acoustic feature encoding was affected both by spectral degradation and language, but language-based effects were generally only present within one or two of the three acoustic conditions. For example, spectral variation encoding weights contained a pre-stimulus cluster (-30-100 ms) peaking at-10 ms, indicating an interaction between vocoded + blurring and Dutch (*F*(2.00,1320.00) = 20.72, *p <* 0.001). In left frontal sensors (Figure 4, Panel B), paired tests showed that vocoded + blurring English weights (Mean = −0.36, SD = 1.36) were significantly more negative than vocoded + blurring Dutch weights (Mean = 0.21, SD = 1.22; Est = 0.57, 95% CI [0.37,0.77], *t*(1,1359.00) = 5.60, *p <* 0.001). Conversely, unprocessed English (Mean = 0.19, SD = 1.14) was slightly but significantly more positively encoded than unprocessed Dutch (Mean = −0.12, SD = 1.12; Est = −0.31, 95% CI [-0.51,-0.11], *t*(1,1359.00) = −3.01, *p* = 0.003). Vocoded English (Mean = 0.09, SD = 1.03) and vocoded Dutch (Mean = −0.04, SD = 1.18) did not differ from one another (*p* = 0.201).

We also identified an early cluster in envelope derivative weights (10-170 ms; Figure 4, Panel C) showing an interaction between vocoded and Dutch, with its peak at about 60 ms across multiple regions of interest. In left frontal sensors (*F*(2.00,1320.00) = 21.63, *p <* 0.001), pairwise contrasts showed that vocoded Dutch weights (Mean = 2.78, SD = 2.33) were significantly more positive than vocoded English weights (Mean = 1.58, SD = 1.60; Est = 1.20, 95% CI [0.95,1.45], *t*(1,1359.00) = 9.33, *p <* 0.001). For unprocessed speech, Dutch (Mean = 1.79, SD = 1.75) and English (Mean = 1.70, SD = 1.74) encoding did not differ, and the contrast between vocoded + blurring Dutch (Mean = 1.68, SD = 1.94) and vocoded + blurring English (Mean = 1.42, SD = 1.59) failed to reach corrected significance (*p* ≥ 0.027).

These effects are broadly in line with others shown in Figure 4; that is, differences between the acoustic feature encoding of comprehensible and incomprehensible speech are limited to particular levels of spectral clarity. By comparison, clearer patterns emerge with respect to spectral degradation (e.g, with spectral flux; Figure 4, Panel A). One exception was for envelope encoding, where we found a cluster (90-160 ms; Figure 4, Panel E) associated with a main effect of Dutch, peaking at 130 ms. Dutch speech was associated with more positive weights (Mean = 0.92, SD = 1.98) compared to English speech (Mean = 0.30, SD = 1.52; *F*(1.00,1325.00) = 60.04, *p <* 0.001). However, this effect only reached cluster-level significance in left frontal sensors (see other regions in Supplementary Material, Figure 10).

Likewise, in envelope derivative weights, there was a late cluster (430-650 ms) associated with a main effect of Dutch that peaked at 600 ms (Figure 4, Panel C). However, the linear mixed effects model at cluster max also revealed an interaction with vocoded (Est = 0.52, 95% CI [0.34,0.70], *t*(1,1360.00) = 5.69, *p <* 0.001). Pairwise tests comparing language showed that only unprocessed English (Mean = −0.52, SD = 1.04) significantly differed from unprocessed Dutch (Mean = 0.00, SD = 0.0.89; Est = 0.52, 95% CI [0.34,0.70], *t*(1,1360.00) = 5.69, *p <* 0.001); there were no significant differences between English and Dutch for either spectrally degraded condition (*p* ≥ 0.101).

In sum, although acoustic feature encoding is modulated by language to a certain extent, we fail to find a robust main effect this is apparent at all levels of spectral clarity.

#### 3.2.2 Linguistic Features

For linguistic feature encoding, we find that language effects are generally more pronounced and sustained in comparison to acoustic feature encoding. However, these effects are nonetheless modulated by spectral clarity. For phoneme onsets (Figure 4, Panel F), for instance, encoding weights contained an early cluster (0-230 ms) with its peak at around 110 ms that indicated an interaction between spectral degradation and Dutch (*F*(2.00,1325.00) = 13.55, *p <* 0.001). Pairwise tests showed that vocoded Dutch weights (Mean = 1.38, SD = 1.87) were significantly more positive than vocoded English weights (Mean = 0.86, SD = 2.16; Est = 0.53, 95% CI [0.23,0.82], *t*(1,1360.00) = 3.48, *p <* 0.001).

Conversely, vocoded + blurring Dutch weights (Mean = 0.67, SD = 1.92) were significantly *less* positive than vocoded + blurring English weights (Mean = 1.15, SD = 1.68; Est = −0.48, 95% CI [-0.78,-0.19], *t*(1,1360.00) = −3.19, *p* = 0.001). The pairwise test between unprocessed Dutch and English did not meet corrected statistical significance (*p* = 0.011).

Staying with phoneme onsets, but in right frontal electrodes (Figure 4, Panel G), we identified a cluster (30-120 ms) with its peak at 90 ms showing an interaction between Dutch and vocoded + blurring (*F*(2.00,1325.00) = 5.23, *p* = 0.005). Unprocessed English encoding (Mean = 0.26, SD = 1.69) had significantly more positive weights than unprocessed Dutch (Mean = −0.77, SD = 2.36; Est = −1.03, 95% CI [-1.40,-0.66], *t*(1,1360.00) = −5.50, *p <* 0.001). Vocoded English (Mean = 0.07, SD = 1.86) was more positive than vocoded Dutch (Mean = −0.55, SD = 2.38; Est = −0.62, 95% CI [-0.98,-0.25], *t*(1,1360.00) = −3.29, *p* = 0.001). However, vocoded + blurring English (Mean = −0.52, SD = 2.23) and vocoded + blurring Dutch (Mean = −0.69, SD = 1.93) did not differ from one another (*p* = 0.355).

In comparison to phoneme onsets and word onsets, syllable onset encoding was generally noisier at the group level, as evidenced by wide confidence intervals of the mean (Supplementary Material, Figure 10). Yet, this feature was associated with a relatively long cluster (250-520 ms) peaking at 400 ms in left frontal sensors (Figure 4, Panel H), revealing an interaction between language and vocoded + blurring (*F*(2.00,1325.00) = 21.44, *p <* 0.001). Whereas unprocessed English (Mean = 0.38, SD = 1.58) did not differ from unprocessed Dutch (Mean = 0.23, SD = 1.99; *p* = 0.329), vocoded English (Mean = 0.37, SD = 1.82) was significantly more positive than vocoded Dutch (Mean = −0.07, SD = 1.45; Est = −0.45, 95% CI [-0.75,-0.14], *t*(1,1360.00) = −2.89, *p* = 0.004). Within vocoded + blurring, the direction of this effect was reversed, with English (Mean = −0.35, SD = 2.04) significantly more negative in comparison to Dutch (Mean = 0.56, SD = 1.63; Est = 0.91, 95% CI [0.61,1.21], *t*(1,1360.00) = 5.91, *p <* 0.001).

Finally, we highlight two clusters associated with word onset encoding. The earlier cluster emerged as a pre-stimulus trend seen across all regions of interest (Supplementary Material, Figure 13), but which only reached cluster-corrected statistical significance in central frontal and left posterior sensors (Figure 4, Panel I). In left posterior electrodes (−150 − −40), this interaction peaked at −90 ms between Dutch and vocoded (*F*(2.00,1325.00) = 9.79, *p <* 0.001). Pairwise contrasts revealed that vocoded English weights (Mean = −1.03, SD = 1.60) were significantly more negative than vocoded Dutch weights (Mean = −0.11, SD = 1.17; Est = 0.92, 95% CI [0.70,1.14], *t*(1,1360.00) = 8.13, *p <* 0.001). Vocoded + blurring English (Mean = −0.42, SD = 1.17) was also more negative than vocoded + blurring Dutch (Mean = 0.00, SD = 1.33; Est = 0.43, 95% CI [0.21,0.65], *t*(1,1360.00) = 3.78, *p <* 0.001). However, the difference between unprocessed English (Mean = −0.42, SD = 1.47) and unprocessed Dutch weights (Mean = −0.19, SD = 1.16) failed to reach corrected significance (*p* = 0.040).

The second cluster for word onset encoding occurred later (260-650 ms) and robustly across all six regions of interest. For left posterior sensors, the effect peaked at about 420 ms as an interaction between vocoded + blurring and Dutch (*F*(2.00,1325.00) = 19.08, *p <* 0.001). Unprocessed English encoding (Mean = 1.43, SD = 1.58) had significantly more positive weights than unprocessed Dutch encoding (Mean = 0.16, SD = 1.09; Est = −1.27, 95% CI [-1.51,-1.03], *t*(1,1360.00) = −10.37, *p <* 0.001), and vocoded English (Mean = 0.95, SD = 1.81) also had significantly more positive weights than vocoded Dutch encoding (Mean = −0.08, SD = 1.30; Est = −1.03, 95% CI [-1.27,-0.79], *t*(1,1360.00) = −8.43, *p <* 0.001). Vocoded + blurring English (Mean = −0.10, SD = 1.41) had less negative weights than vocoded + blurring Dutch (Mean = −0.34, SD = 1.27), but this contrast failed to reach corrected significance (*p* = 0.043).

Taken together, although we do find ample evidence for the modulation of feature processing by language, these effects are subject to change in different acoustic contexts. The linear mixed effects modelling shows that–whether for acoustic or linguistic features–there are no clear and consistent effects of language on neural speech encoding.

## 4 Linguistic Feature Encoding Does Not Scale with Self-Reported Comprehension

We observed robust interactions with language during linguistic feature encoding–for instance, for word onsets (Figure 4, Panel I). Although the magnitude of word onset encoding is greater for unprocessed and vocoded English in comparison to their corresponding Dutch conditions, word onset encoding for the vocoded + blurring condition is indistinguishable during English and Dutch listening. If we assume that effects of language reflect comprehension, then we could hypothesise that participants simply did not understand this most spectrally degraded English condition. However, we also collected self-reported intelligibility ratings showing that many participants achieved at least moderate understanding of vocoded + blurring English (Figure 2, Panel A). This subjective measure is corroborated by objectively measured response times to auditory targets in the repeated phrase detection task, which correlate to reported comprehension (rho = 0.52, *p* = 0.001; Figure 2, Panel B).

To further explore the relationship between comprehension of vocoded + blurring English and linguistic feature encoding, we identified the minimum and maximum values of Phoneme Onset and Word Onset weights within participant for each region of interest. This resulted in 24 dependent measures (2 measures by 2 features by 6 regions), for each of which we assessed the correlation with intelligibility ratings. Even at a liberal, uncorrected threshold of significance (*p* = 0.05), there were only two neural predictors of comprehension. These were the maximum value of Word Onset weights in left posterior sensors, Rho = −0.36 (*p* = 0.027, 95% CI [−0.62, −0.03]), and the maximum value of Phoneme Onset weights in central frontal sensors, Rho = 0.44 (*p* = 0.006, 95% CI [0.13, 0.68]). Note that neither of these associations would survive Bonferroni correction (i.e., *p* = 0.002 would be required for corrected significance).

In addition, we also observed a lack of *specificity* for linguistic feature encoding. This can be seen in Figure 5, where we display results of participants split in tertiles based on self-reported comprehension of vocoded + blurring English. Panel F, for instance, illustrates how some Dutch listening conditions elicit relatively large phoneme encoding weights, particularly for participants who report the best understanding of vocoded + blurring English. Conversely, those same participants showed encoding of word onsets only for unprocessed English, despite finding degraded English conditions highly intelligible (Figure 5, Panel C). Taken together, we find poor reliability of linguistic feature weights for predicting comprehension; that is, neither word onset nor phoneme onset encoding provide a direct or robust neural indicator of comprehension for individual participants across all listening conditions.

**Figure 5:**
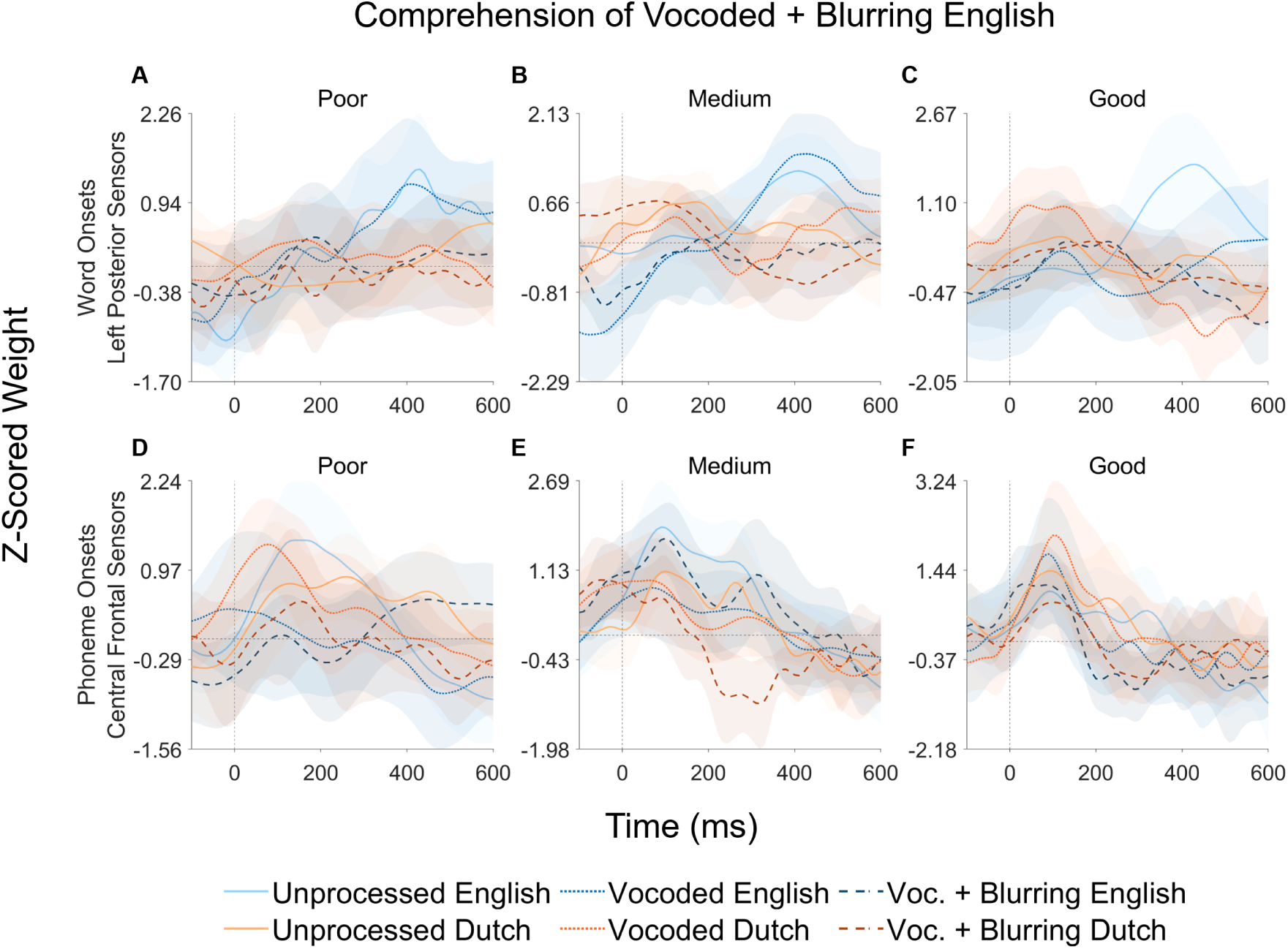
Z-scored encoding model weights across experimental conditions for linguistic features that correlated with self-reported comprehension of vocoded + blurring English. Data are grouped into tertiles by understanding level for descriptive purposes: Poor (n = 12), medium (n = 15), and good (n = 11). Traces and shaded regions indicate the group mean (averaged over sensors) and 95% confidence intervals of the mean.

## 5 Flexible Neural Encoding Predicts Comprehension of Spectrally Degraded English

Given that encoding of single features within single conditions was not specifically or reliably associated with comprehension, we applied a data-driven method to combine multiple measures of neural encoding *across* conditions to predict understanding (see Methods for details). Briefly, we calculated neural contrasts by subtracting the encoding weights of one condition (the comparison; e.g., vocoded + blurring English) from another (the reference; e.g., unprocessed English). We identified features and time points where neural contrasts were correlated to comprehension, and which survived cluster-based permutation tests. This yielded a subset of 30 neural contrasts all showing moderate bivariate associations with comprehension of a magnitude similar to the two linguistic features reported in the previous section (range rho 0.40 to 0.57; Supplementary Material, Table 12). These 30 neural contrasts were then submitted to an exploratory principal component analysis (PCA) to determine whether mapping them together onto a latent component would improve predictive power, measured as Spearman’s rho with comprehension. To ensure robustness, all PCA analyses were performed using repeated resampling with 50% of participants held out for testing.

We empirically determined that the maximum rho could be reached by combining as few as ten neural contrasts (Figure 6, Panel A). We then selected which ten neural contrasts to include based on the frequency of their contributions to the top 5%, or most predictive, of possible PCA solutions (range predictive rho 0.83 to 0.88; Figures 7-8). Importantly, although we highlight one particular solution here, comparable performance can be achieved using multiple alternative combinations of neural contrasts: There were 18 distinct neural contrasts that appeared at least once in the best combinations (Table 2). The encoding weights associated with the ten neural contrasts included in the final PCA are shown in Figure 8, Panel B. The weights for all 18 neural contrasts appearing in the top 5% of possible PCA solutions are provided in the Supplementary Material, Figures 13-16.

**Figure 6:**
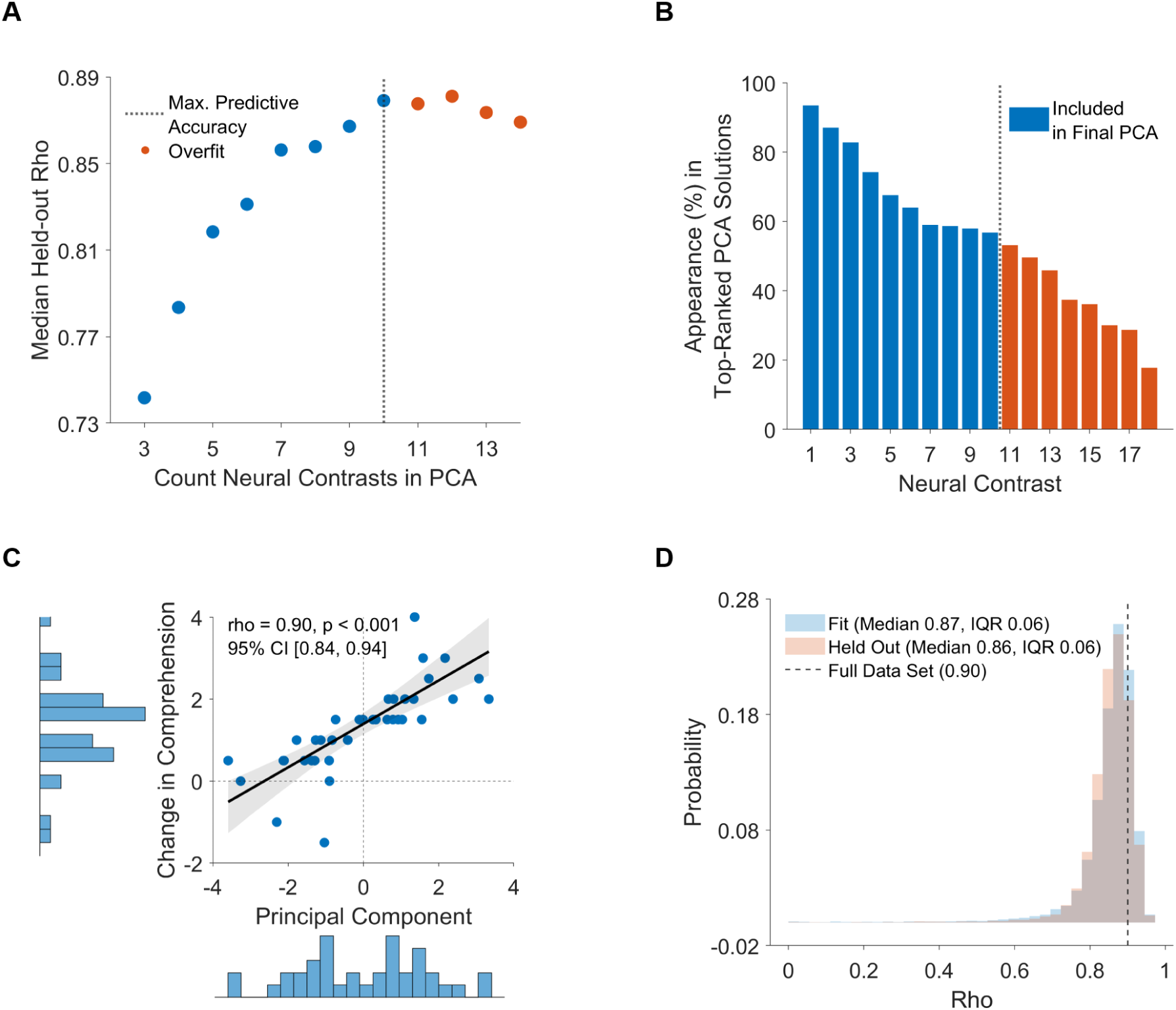
Panel A: Relationship between the number of neural contrasts that contribute to the principal component analysis (PCA) and median held-out rho between the first principal component and comprehension of vocoded + blurring English. 50% of participants (*n* = 19) are held out for testing rho across repeated random resampling with 10,000 iterations. Panel B: neural contrasts ordered by the frequency of their contribution to the top 5%-ranked PCA solutions (i.e., 2,188 of 43,758 possible combinations of neural contrasts) where the first principal component predicted comprehension of vocoded + blurring English. Details regarding each neural contrast are given in Table 2. Panel C: Distributions and bivariate association between the first principal component of the final PCA containing 10 neural contrasts and comprehension of vocoded + blurring English, calculated using the full data set (*n* = 38). Lines and shaded regions indicate the least-squares linear fit with bootstrap 95% confidence intervals. Panel D: Histograms depicting the distributions of rho between the first principal component of the final PCA with 10 neural contrasts and comprehension of vocoded + blurring English for fit and held-out data across repeated random resampling.

**Figure 7:**
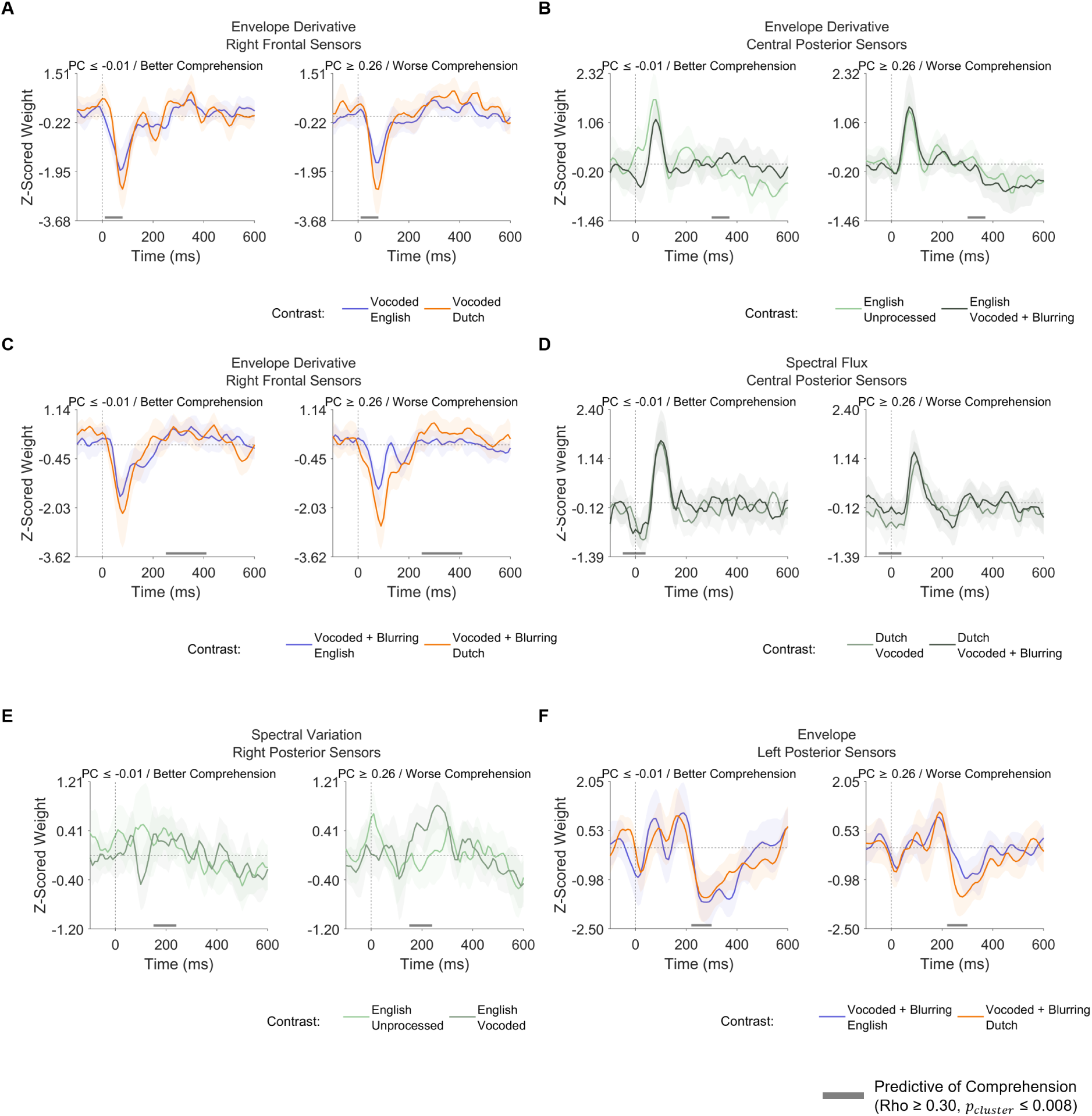
Encoding weights associated with the neural contrasts contributing most frequently to the top 5%-ranked PCA solutions, descriptively grouped by median-split principal component score. The panels are ordered by the frequency of each neural contrast’s contribution to the top solutions. Grey bars indicate temporal regions where a correlation with comprehension of vocoded + blurring English for that neural contrast was found according to cluster-based permutation testing (5,000 iterations). Traces and shaded regions indicate the group mean (averaged over sensors) and 95% confidence intervals of the mean.

**Figure 8:**
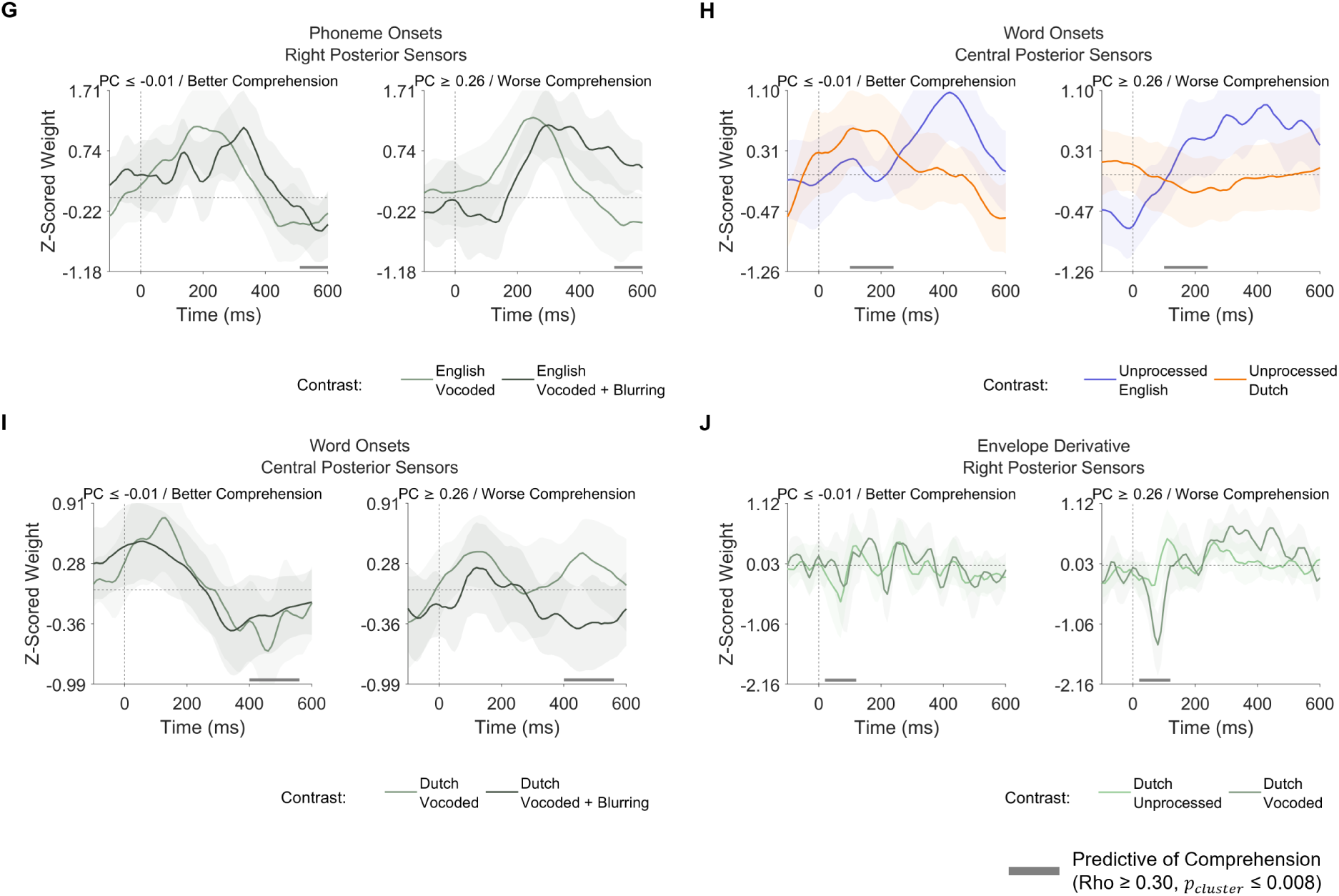
Encoding weights associated with the neural contrasts contributing most frequently to the top 5%-ranked PCA solutions, descriptively grouped by median-split principal component score. The panels are ordered by the frequency of each neural contrast’s contribution to the top solutions. Grey bars indicate temporal regions where a correlation with comprehension of vocoded + blurring English for that neural contrast was found according to cluster-based permutation testing (5,000 iterations). Traces and shaded regions indicate the group mean (averaged over sensors) and 95% confidence intervals of the mean.

**Table 2:** Neural contrasts individually predictive of comprehension of vocoded + blurring English. Ranking and inclusion in the final PCA containing 10 neural contrasts is determined by the frequency of its contribution to the top 5%-ranked PCA solutions (i.e., 2,188 of 43,758 possible combinations of neural contrasts) where the first principal component predicted comprehension of vocoded + blurring English. The significance of clusters is identified using mass permutation tests with 5000 iterations. Abbreviations: CF Central Frontal; LF Left Frontal; RF Right Frontal; CP Central Posterior; LP Left Posterior; RP Right Posterior. The threshold of significance for multiple comparisons over six regions of interest is set at *p_cluster_ ≤* 0.008.

| Rank | In Final PCA | Feature | Cluster |  |  | Time (ms) |  |  |  | Rho |  |  |
| --- | --- | --- | --- | --- | --- | --- | --- | --- | --- | --- | --- | --- |
| | | | ROI | Reference | Comparison | Start | Max | End | Length | Sum | Max | $p_{cluster}$ |
| 1 | Yes | EnvDeriv | RF | Voc. En. | Voc. Du. | 10 | 40 | 80 | 70 | 1.74 | 0.52 | < 0.0001 |
| 2 | Yes | EnvDeriv | CP | Unproc. En. | Voc. Blur. En. | 300 | 350 | 370 | 70 | 1.63 | 0.5 | 0.0020 |
| 3 | Yes | EnvDeriv | RF | Voc. Blur. En. | Voc. Blur. Du. | 250 | 280 | 410 | 160 | 4.85 | 0.48 | < 0.0001 |
| 4 | Yes | SpectralFlux | CP | Voc. Du. | Voc. Blur. Du. | -50 | -10 | 40 | 90 | 2.73 | 0.57 | < 0.0001 |
| 5 | Yes | SpectralVariation | RP | Unproc. En. | Voc. En. | 150 | 190 | 240 | 90 | 2.24 | 0.45 | < 0.0001 |
| 6 | Yes | Envelope | LP | Voc. Blur. En. | Voc. Blur. Du. | 220 | 260 | 300 | 80 | 1.85 | 0.41 | < 0.0001 |
| 7 | Yes | PhonemeOnsets | RP | Voc. En. | Voc. Blur. En. | 510 | 580 | 600 | 90 | 2.42 | 0.45 | < 0.0001 |
| 8 | Yes | WordOnsets | CP | Unproc. En. | Unproc. Du. | 100 | 160 | 240 | 140 | 4.06 | 0.43 | < 0.0001 |
| 9 | Yes | WordOnsets | CP | Voc. Du. | Voc. Blur. Du. | 400 | 450 | 560 | 160 | 4.75 | 0.47 | < 0.0001 |
| 10 | Yes | EnvDeriv | RP | Unproc. Du. | Voc. Du. | 20 | 70 | 120 | 100 | 2.94 | 0.47 | < 0.0001 |
| 11 | No | SpectralFlux | RP | Unproc. Du. | Voc. Blur. Du. | 60 | 90 | 120 | 60 | 1.09 | 0.41 | 0.0060 |
| 12 | No | SpectralFlux | RP | Voc. Blur. En. | Voc. Blur. Du. | 50 | 90 | 120 | 70 | 1.65 | 0.48 | < 0.0001 |
| 13 | No | SpectralVariation | LP | Voc. En. | Voc. Blur. En. | 240 | 280 | 320 | 80 | 2.04 | 0.5 | < 0.0001 |
| 14 | No | WordOnsets | CF | Unproc. Du. | Voc. Du. | 350 | 440 | 500 | 150 | 4.74 | 0.44 | < 0.0001 |
| 15 | No | Envelope | RP | Unproc. En. | Unproc. Du. | 220 | 290 | 320 | 100 | 2.66 | 0.43 | < 0.0001 |
| 16 | No | Envelope | LF | Voc. Blur. En. | Voc. Blur. Du. | 350 | 380 | 410 | 60 | 1.17 | 0.44 | 0.0030 |
| 17 | No | SyllableOnsets | CF | Voc. Blur. En. | Voc. Blur. Du. | 500 | 570 | 600 | 100 | 2.84 | 0.45 | < 0.0001 |
| 18 | No | EnvDeriv | LF | Voc. En. | Voc. Blur. En. | -50 | -10 | 20 | 70 | 1.67 | 0.48 | 0.0020 |

Of the ten neural contrasts that we retained in the final PCA, seven neural contrasts involved acoustic features and three involved linguistic features. When run on the complete data set, the first principal component eigenvalue was 2.84 and explained 28.5% of variance. It predicted comprehension with rho = 0.90 (Figure 6, Panel C). We inspected the coefficients or loadings, which ranged from −0.32 to 0.34. The unsigned Mean value was 0.32 (SD = 0.02). Thus, the neural contrasts were roughly equal in importance and no single predictor dominated the principal component. When the PCA was re-fit across 10,000 random iterations with 50% held-out data, median held-out rho only slightly decreased to 0.86 (IQR = 0.06; Figure 6, Panel D). The consistent performance on split-half data shows that the latent component we identified is stable and that its relationship to comprehension generalises across individuals within this study.

Notably, only four of the ten neural contrasts in the final PCA directly involved the vocoded + blurring English condition. In fact, the neural contrast that contributed most consistently was English − Dutch contrast within *vocoded* speech for envelope derivative encoding, which occurred 20-60 ms post-stimulus onset in right frontal electrodes (Figure 7, Panel A). When we re-ran PCA using only neural contrasts that directly contrasted vocoded + blurring English with another condition (e.g., vocoded + blurring Dutch), median held-out rho dropped to = 0.70 (IQR = 0.13). In other words, using the neural response to unrelated experimental conditions improves our ability to estimate individual comprehension of spectrally degraded speech. This suggests that the full PCA captures a listener *profile* or traits that are consistently associated with better understanding, but does not necessarily indicate the specific mechanisms directly underpinning this advantage.

## 6 Discussion

Neural encoding models provide insights into the mechanisms underlying perception and cognition. However, most studies implicitly assume that measuring typical responses at the group level will enable the detection of differences in neural encoding between participants that are linked to comprehension. We used a crossed study design to assess the impact of linguistic comprehension while varying acoustic sensory processing. Our results show that neural encoding for single listening conditions may offer a limited or even misleading view. Namely, when comparing the neural encoding of speech features at the group level, we found that no single acoustic or linguistic feature consistently signalled successful understanding; instead, we observed disparate patterns in the magnitude and direction of encoding differences across listening conditions which did not predict speech comprehension.

We further showed that encoding differences or contrasts between listening conditions are systematically related to comprehension Combining both acoustic and linguistic features in a data-driven manner using PCA we were able to predict comprehension of spectrally degraded speech in individual listeners with good out-of-sample accuracy. This finding challenges other proposals that linguistic feature encoding is reliably, or exclusively, linked to comprehension.

Instead, our findings provide evidence that flexibility in neural encoding of speech is an informative trait that can reveal changes in perceptual and cognitive processing strategies under challenging listening conditions. Moreover, we show that this trait appears to vary across individuals in a way that predicts comprehension with a high level of accuracy. If the current results are born out in future work, this method may provide a neural correlate of individual differences in speech perception that are linked to behaviour, such as cue weighting or word recognition. It could also potentially serve as an objective measure for the diagnosis of hearing or comprehension difficulties.

Apart from applying PCA to neural contrasts across conditions, another important aspect of this study is that we assess multiple acoustic features beyond the amplitude envelope, including features that characterise change in spectral information over time (MacIntyre et al., 2026). We algorithmically compared encoders built from various combinations of acoustic features and found that, despite its ubiquitous use in speech perception research, the broadband envelope is less effective at explaining the neural response to speech than spectral flux, spectral variation, or the envelope first derivative. In other words, the amplitude envelope is an ineffective stand-in for speech acoustics in general. Furthermore, and contrary to some recent work comparing acoustic to linguistic features (Karunathilake et al., 2023; Gillis et al., 2023), we found effects of language on acoustic encoder weights. That said, these effects were relatively diffuse and short-lived. Had we stopped there, the current study would have agreed with previous work proposing that the neural encoding of acoustic features primarily reflects low-level sensory processing. However, in our data-driven analysis of individual differences in neural encoding that are linked to comprehension, seven of the ten best explanatory neural measures were derived from acoustic feature encoding weights. Thus, we conclude that neural encoding of speech acoustics is modulated by both sensory and cognitive influences during speech understanding which are linked to listening outcomes.

### 6.1 Neural encoding of linguistic events dissipates under spectral degradation, despite comprehension

Numerous studies have found robust encoding of word onsets and other lexical features with weights peaking at approximately 400 ms post-stimulus onset (e.g., Broderick et al., 2021; Heilbron et al., 2022; Broderick et al., 2022; Gillis et al., 2023; Puffay et al., 2023; Chalehchaleh et al., 2025). This in turn builds on decades of work with event-related potential methods that establish the canonical n400 response as an electrophysiological landmark associated with lexical processing (Kutas and Federmeier, 2011). When grouping our participants according to their comprehension of vocoded + blurring English, we observed that word onsets are only consistently encoded for unprocessed English, but not for lower clarity conditions (Figure 5, Panels A-C). Specifically, for the spectrally degraded, but equivalently intelligible vocoded English condition, ‘poor’ and ‘medium’ comprehenders show the anticipated 400 ms peak; however, this response is visibly attenuated for ‘good’ comprehenders. Finally, in the vocoded + blurring English condition, no evidence for word onset encoding emerges for any group. We speculate that this unexpected pattern may relate to shifting processing strategies under varying acoustic conditions. Vocoding removes spectral detail from the speech signal, whilst preserving temporal structure, including the envelope. Using PCA, we found that changes to word onset encoding are systematically related to concomitant shifts in acoustic feature encoding. Hence, it is possible that diminished word onset encoding mirrors low-level mechanisms, such as the down-weighting of less informative acoustic cues (i.e., those of a spectral rather than temporal nature).

A higher level—but not mutually exclusive—interpretation is proposed in the literature on incremental word recognition. Spoken language comprehension requires listeners to map the continuous speech signal onto lexical representations in real time (Christiansen and Chater, 2016). Prominent theories of how this is achieved propose that listeners rapidly assess competing lexical candidates that are compatible with the acoustic input received so far (Marslen-Wilson, 1984; Weber and Scharenborg, 2012). For example, a word that begins with’ca-’ could conclude as’cat’,’cap,’ or’cab’, but not’cope’ or’coffee’. Once the listener hears’-b’, then’cat’ and’cap’ can also be excluded. Although the details of how listeners reject or filter out incompatible words are debated (Jusczyk and Luce, 2002; Magnuson et al., 2007; Vitevitch and Luce, 2016), this framework has received abundant empirical support from listening experiments under controlled, acoustically ideal conditions with neurotypical listeners (Eberhard et al., 1995; Jusczyk and Luce, 2002; Kapnoula et al., 2017).

Interestingly, data from clinical populations indicate that audiological or developmental differences may produce divergent word recognition trajectories. Cochlear implant (CI) users, for instance, show delayed word recognition and reduced competition effects for words with the same initial sound (McMurray et al., 2017; Colby et al., 2024; McMurray et al., 2024). This may suggest that CI users wait to acquire further sensory information before determining lexical identity, an explanation that was supported by another experiment involving typically hearing listeners presented with vocoded speech, which simulates the limited spectral resolution conveyed by CI (McMurray et al., 2017). This “wait-and-see” strategy could help to explain the absence of a time-locked word onset response for vocoded + blurring English in the current study, even in listeners who, like many CI users, are nonetheless able to achieve good understanding of noisy or ambiguous sensory input. Taken together, word onset encoding may be indicative of successful speech understanding under ideal listening conditions in typically hearing listeners, but it is unlikely to serve as a universal marker of comprehension.

### 6.2 Flexible feature encoding explains individual differences in listening outcomes

Our data-driven analysis shows that there are many neural contrasts that are weakly to moderately associated with comprehension of spectrally degraded English. Using PCA allowed us to combine these neural contrasts and harness their shared variance to produce a composite measure that strongly predicts individual understanding in a held-out sample. Surprisingly, encoding contrasts unrelated to vocoded + blurring English were nonetheless key contributors to this aggregate measure. These include, for example, contrasts generated by comparing different levels of clarity within Dutch, or English against Dutch within the unprocessed condition. Both better and worse comprehenders are alike in behaviour when perceiving these forms of speech: They can all understand clear English and not Dutch irrespective of acoustic clarity. Yet, their neural processing systematically differs for these conditions, and such differences are consistently linked with self-rated comprehension in other listening contexts. Such “variation in process despite equivalent end-state performance” underscores the promise of individualised approaches in the analysis of neural data from speech perception studies (McMurray et al., 2023). Moreover, our results challenge the notion of an “average” listener; it is possible that individuals naturally vary across multiple dimensions in the listening strategies they use in diverse speech processing conditions, yet they are capable of achieving understanding under most circumstances (Yu and Zellou, 2019; Muegge et al., 2026). On the other hand, characterising individual listeners may also provide important insights into speech and hearing disorders, such as identifying which listeners are likely struggle in particular acoustic environments, or experience difficulties in adapting to a new assistive hearing device (Rotman et al., 2020; Cherri et al., 2024). If certain forms of listening flexibility are beneficial for resilient speech perception, then neural measures of how these are recruited in different listening conditions may improve our understanding of developmental or acquired disorders such as aphasia (e.g., by helping to pinpoint the level of processing at which comprehension problems arise; Kries et al., 2024), or suggest new forms of neural intervention.

### 6.3 Limitations

Despite these successes, the current study has limitations which should also be considered. Firstly, we did not collect objective measures of speech comprehension and rely on self-reported ratings in the analysis of individual differences. We opted not to include additional speech tests (e.g., word identification, or comprehension questions) as these tasks would not be applicable to the Dutch conditions, and would also add considerable to the duration of our experimental sessions. However, substantial research into the relationship between objective and subjective measures of comprehension has shown that they closely correspond in young, typically hearing listeners (e.g., Cox et al., 1991; Cienkowski and Speaks, 2000; Kuk et al., 2023; Wiggins et al., 2025). Moreover, we also found high correlations between participants’ response times to auditory targets in the repeated phrase detection task and their comprehension ratings, corroborating their self-reports. Finally, the neural principal component was highly predictive of comprehension for random samples of held-out participants. If self-reported ratings were unreliable or inconsistent, we wouldn’t expect to observe such robust generalisation. Although further work would benefit from including a variety of measures of speech understanding, we are therefore confident that self-reported comprehension is a meaningful and useful measure in the present study.

Another limitation is that we have not provided an in-depth analysis of the neural contrasts which differ between listening conditions; for instance, we do not speculate why better comprehenders tend to have more negative vocoded + blurring English envelope weights, or why poorer comprehenders have more positive vocoded Dutch word onset weights. Our PCA (Shlens, 2014) showed that, the most predictive neural composite score combined ten different contrasts with approximately equal loadings. Hence, although we found the first principal component to have strong predictive value, the precise neural mechanisms underlying successful comprehension and their implications for speech processing theory are not yet fully determined. Future experiments incorporating different listening conditions (e.g., using noisy or reverberant speech) may be of use here, as well as testing more controlled stimuli that systematically vary on specific acoustic or linguistic dimensions, or parametrically varying different combinations of specific neural predictors to infer their importance to the PCA solution.

Finally, the current results demonstrate the importance of investigating diverse characteristics of speech, especially different acoustic features. Although we empirically determined which low-level stimulus properties to include in the encoders, we were not exhaustive in our testing different linguistic features. For example, we did not find lexical surprisal to improve encoder accuracy. However, there are multiple ways of estimating information theoretic measures for speech (Gwilliams and Davis, 2022; Slaats and Martin, 2025) and it is possible that alternative techniques (e.g., using n-gram rather than GPT-based methods; Boeve and Bogaerts, 2025) would produce different results. Given other studies have reported robust surprisal encoding (e.g., Weissbart et al., 2020; Gillis et al., 2021; Puffay et al., 2023; Kries et al., 2024), the present results do not argue against the importance of lexical surprisal or other high-level features during speech understanding.

### 6.4 Conclusion

Listeners overcome considerable variation to comprehend speech across circumstances, speakers, and acoustic environments. How this resilience is achieved remains a key question for auditory cognitive neuroscience. As we have shown, encoding of speech features varies between individuals and, depending on the availability of acoustic cues and linguistic access, this variation can be used to index listening outcomes. We here demonstrate that condition-specific changes in neural encoding, rather than neural encoding per-se, provides an informative and generalisable measure of whether individual listeners can understand speech in challenging conditions involving presentation of degraded speech. These findings point to the utility of precision approaches to the analysis of neural data and suggest potential for improving our understanding of speech processing differences, thereby paving the way for neural diagnosis and intervention where there is need.

## 7 Data and Code Availability

The data and analysis scripts that support the findings of this study are openly available in https://osf.io/2w4ey/.

## 8 Author Contributions

A.D.M.: Conceptualization, Methodology, Programming, Formal analysis, Investigation, Writing - Original Draft, Visualization, Funding acquisition; TG: Conceptualization, Investigation, Writing - Original Draft, Funding acquisition. MHD: Conceptualization, Investigation, Writing - Original Draft, Funding acquisition.

## 9 Funding

A.D.M. was supported by a Leverhulme Trust (ECF-2023-539) and Isaac Newton Trust Early Career Fellowship Award.

T.G. was funded by Medical Research Council UK (MR/T03095X/1). M.H.D. was supported by UK Medical Research Council funding of the MRC Cognition and Brain Sciences Unit (MC_UU_00030/6).

## 10 Declaration of Competing Interests

The authors report no conflicts of interest.

## 11 Supplementary Material

Additional data and analyses supporting this study are included within the article as a Supplementary Material.

## Notes

https://osf.io/2w4ey/

### Competing Interest Statement

The authors have declared no competing interest.

### Summary of Updates

Revised for minor clarifications in main text and figure captions.

