## Supplementary Material for "Flexible neural encoding predicts the comprehension of degraded speech"

### 1 Model Selection

#### 1.1 Stepwise comparison of held-out r and mean-normalised root mean square error

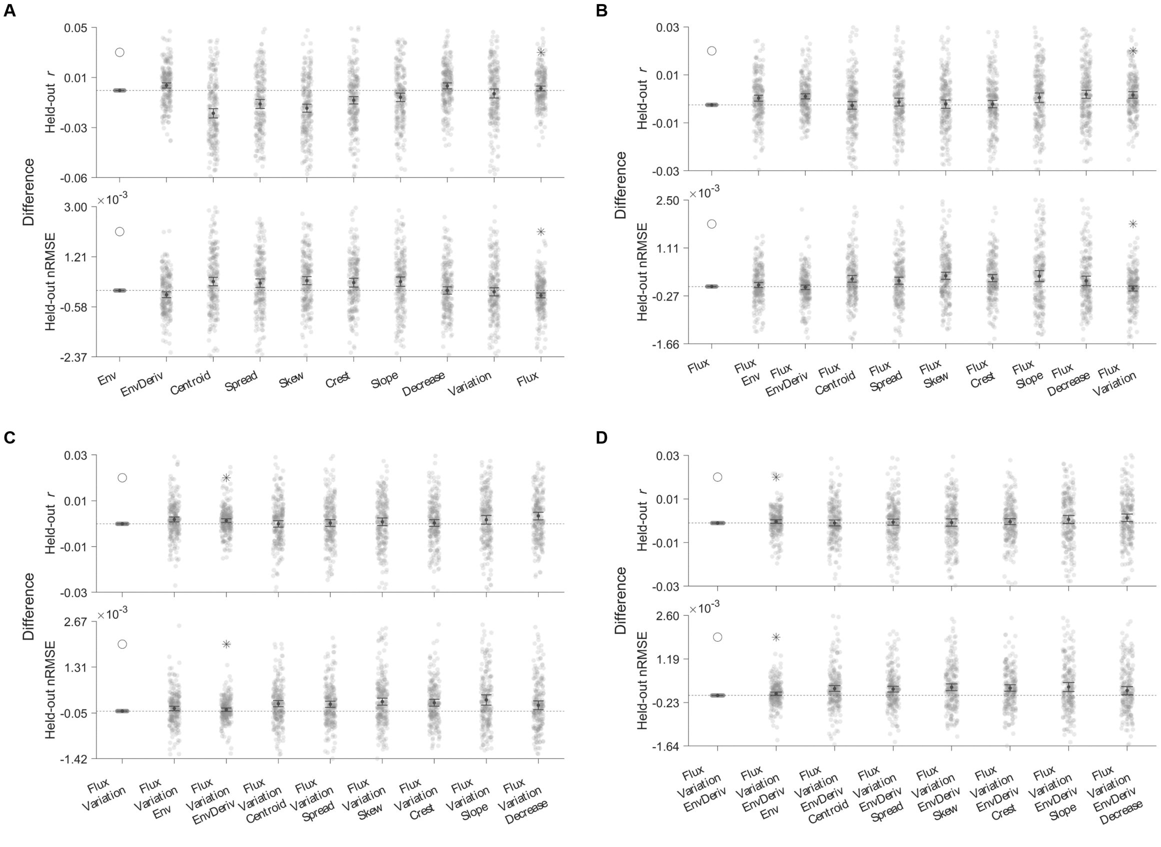

Figure 1: Results of the stepwise model selection procedure for acoustic features. Markers depict the group mean with bootstrap 95% confidence intervals of the mean (5,000 iterations). For each step, the winning model (∗) is determined by its improvement in held out $r$ compared to the baseline model (○) with minimal increase to out-of-sample mean-normalised root mean square error (nRMSE), aggregated across experimental conditions and sensor regions of interest. At the close of each step, the winning feature is included in the subsequent phase baseline model. Panel A: Model 1; Panel B: Model 2. Panel C: Model 3. Panel D: Model 4.

#### 2 Encoding model weights by sensor region of interest

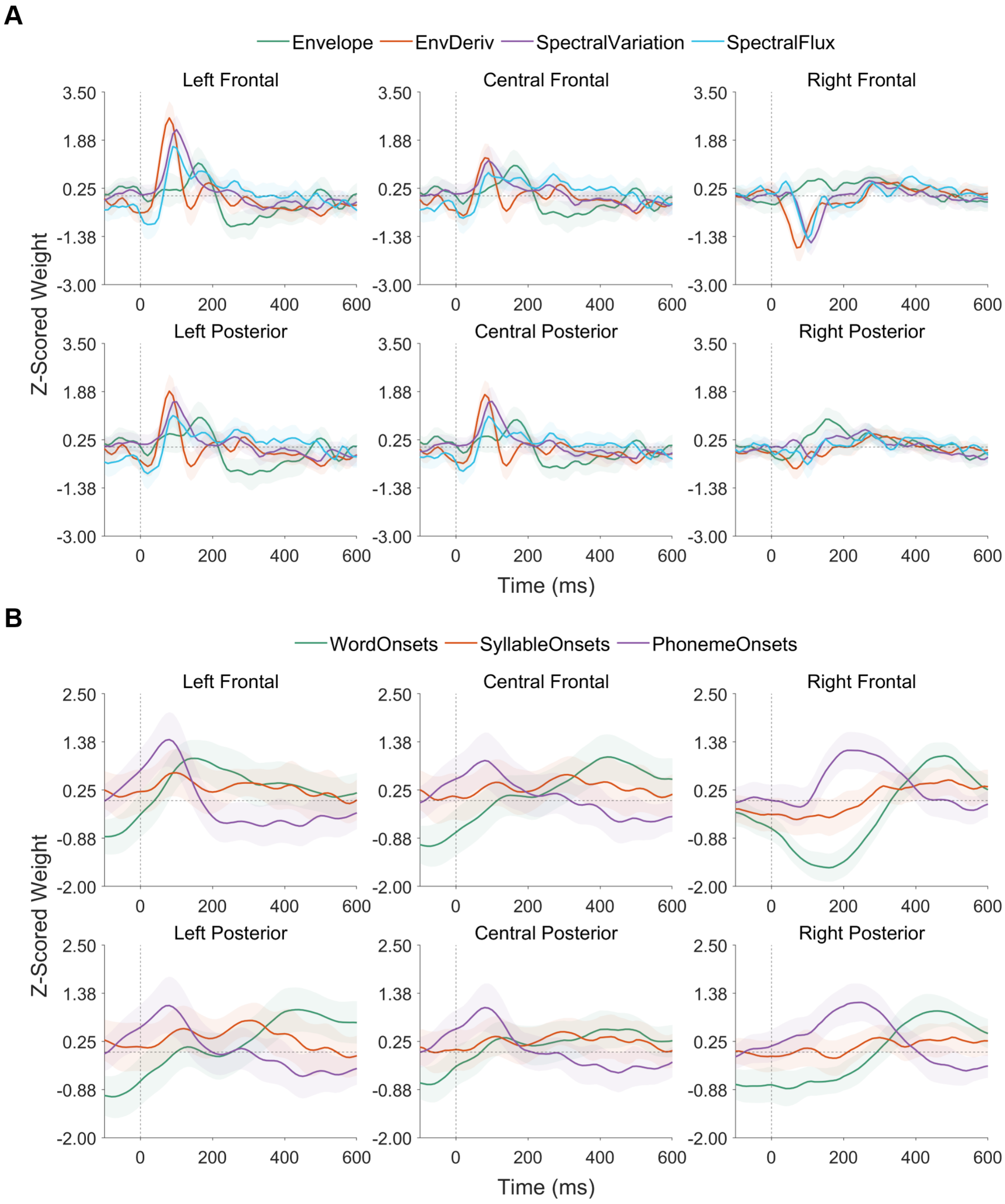


Figure 2: Acoustic (Panel A) and linguistic (Panel B) encoding model weights for Vocoded English by sensor region of interest. Traces represent the group-level mean across sensors within each region of interest; the shaded areas are bootstrap 95% confidence intervals of the mean.

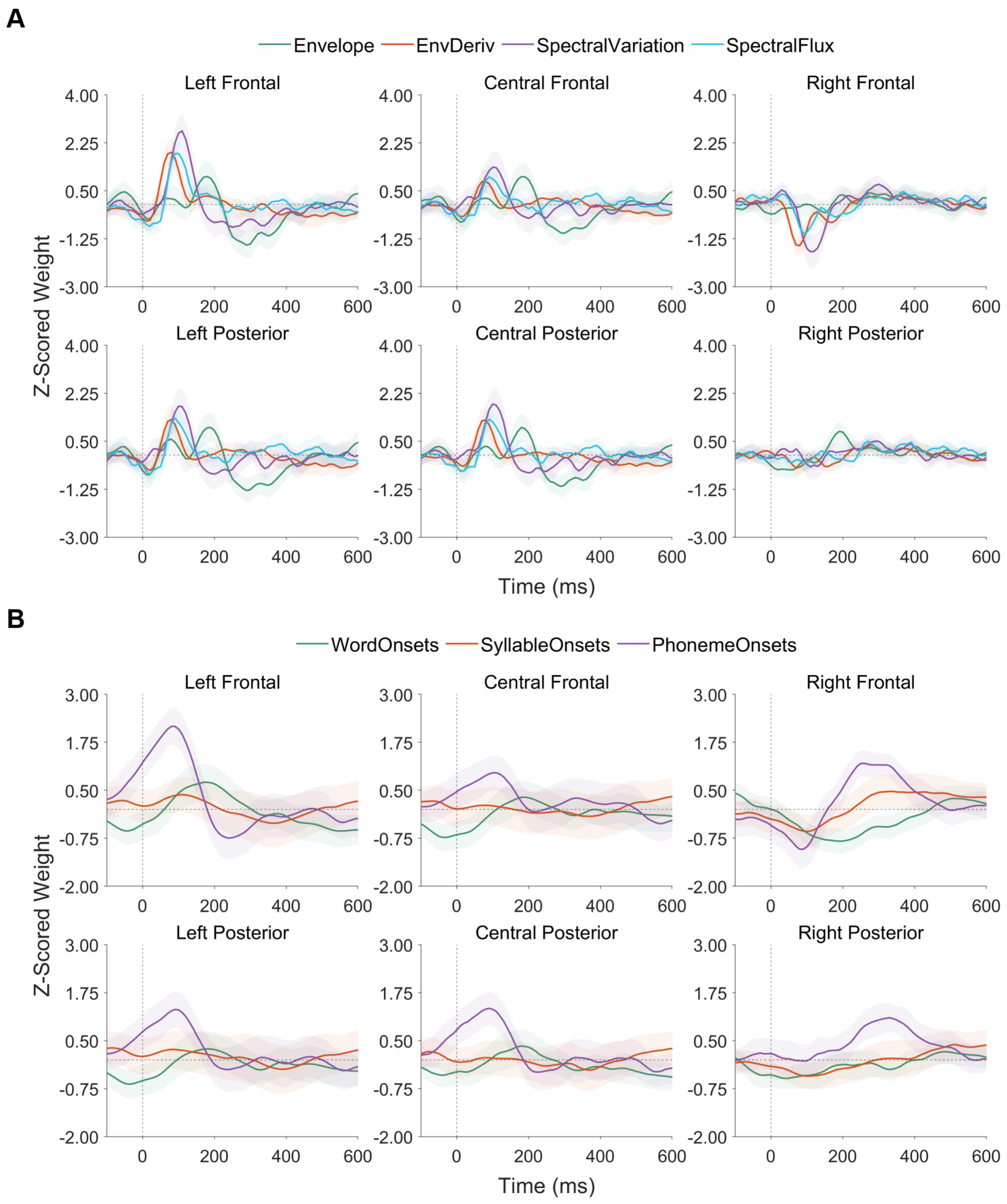


Figure 3: Acoustic (Panel A) and linguistic (Panel B) encoding model weights for Vocoded + Blurring English by sensor region of interest. Traces represent the group-level mean across sensors within each region of interest; the shaded areas are bootstrap 95% confidence intervals of the mean.

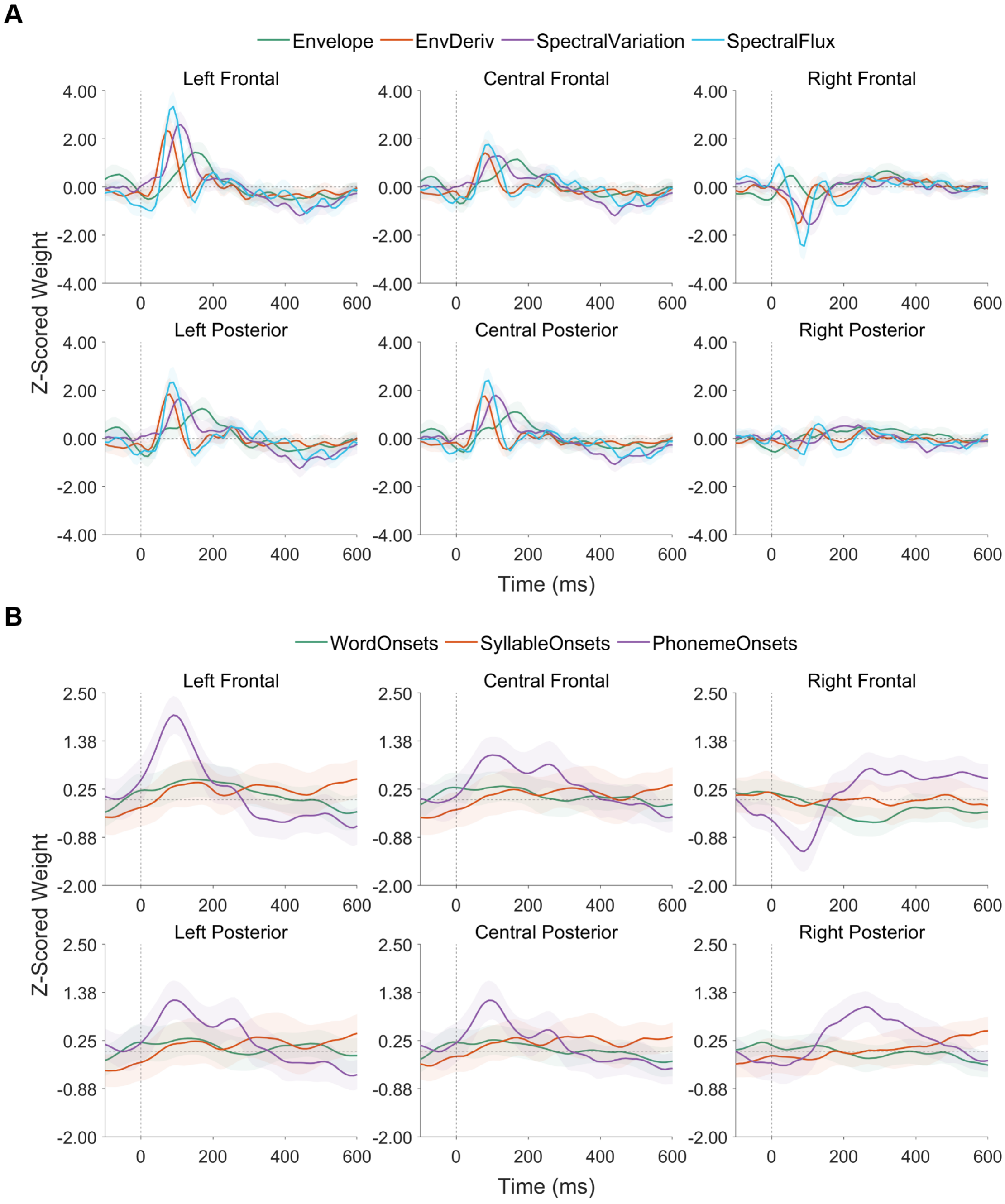


Figure 4: Acoustic (Panel A) and linguistic (Panel B) encoding model weights for Unprocessed Dutch by sensor region of interest. Traces represent the group-level mean across sensors within each region of interest; the shaded areas are bootstrap 95% confidence intervals of the mean.

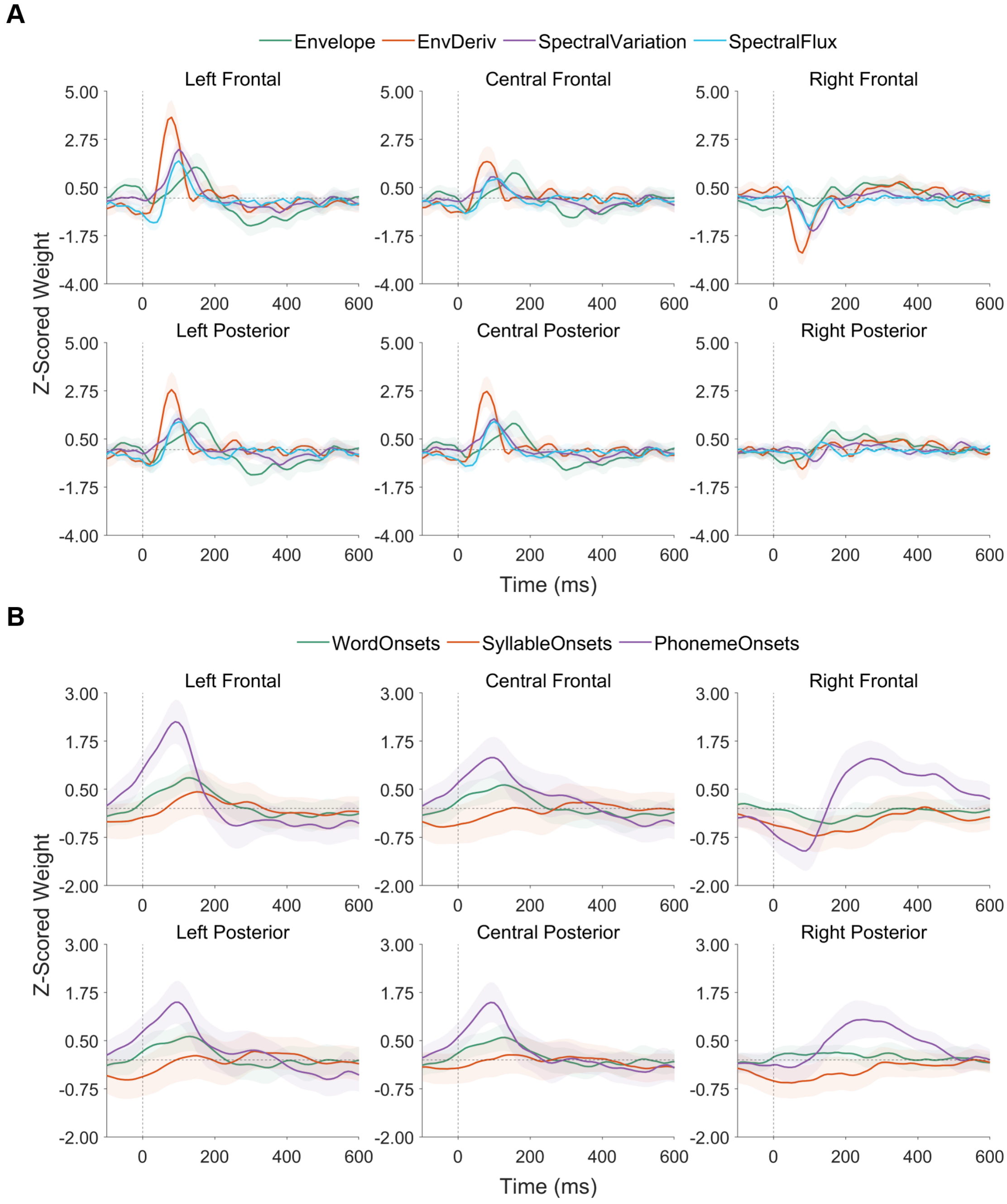


Figure 5: Acoustic (Panel A) and linguistic (Panel B) encoding model weights for Vocoded Dutch by sensor region of interest. Traces represent the group-level mean across sensors within each region of interest; the shaded areas are bootstrap 95% confidence intervals of the mean.

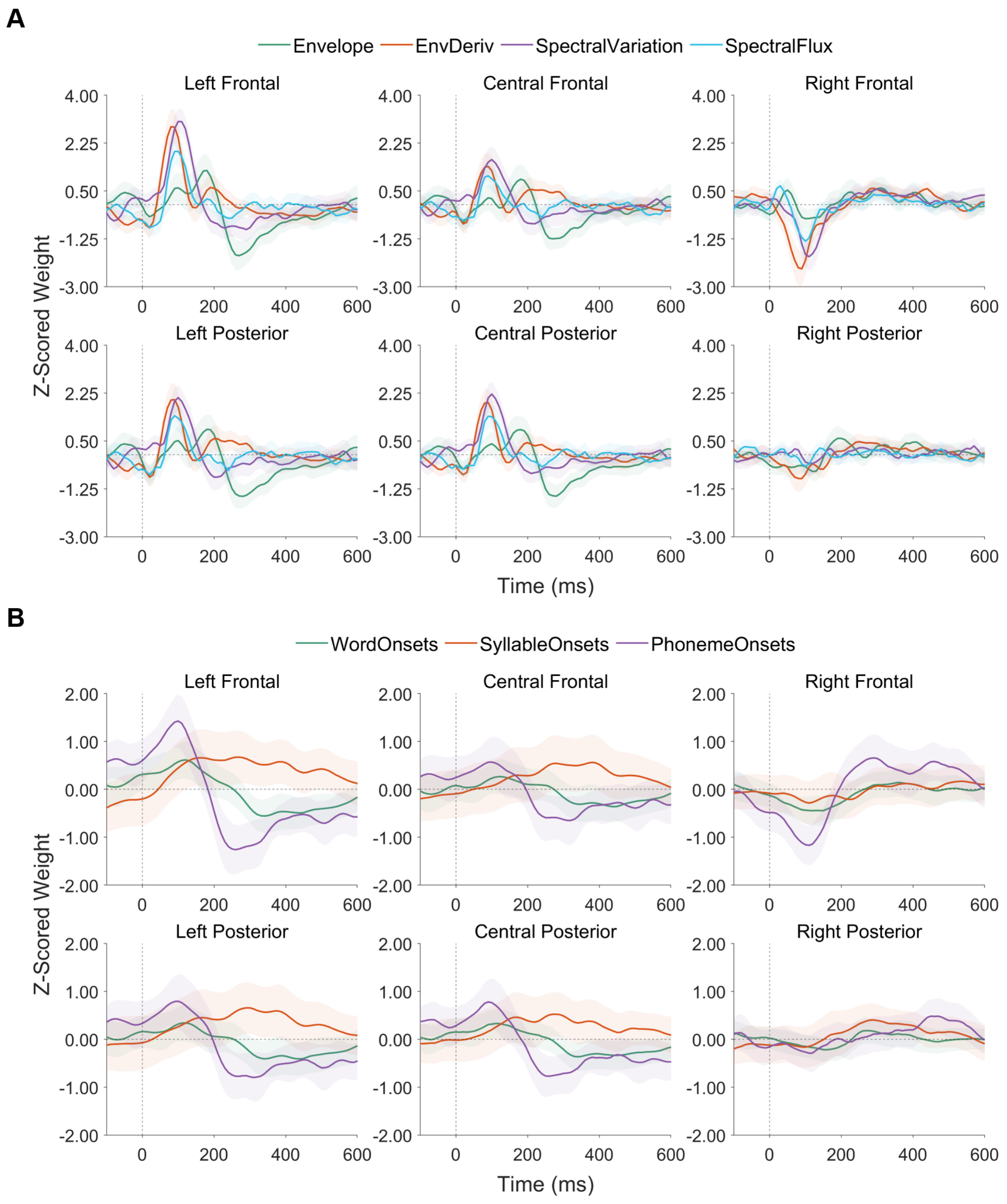


Figure 6: Acoustic (Panel A) and linguistic (Panel B) encoding model weights for Vocoded + Blurring Dutch by sensor region of interest. Traces represent the group-level mean across sensors within each region of interest; the shaded areas are bootstrap 95% confidence intervals of the mean.

#### 3 Latency of peak encoding weights within listener for unprocessed English

The stepwise model selection procedure for acoustic features is intended to establish the unique contributions of each included feature to explain the neural response to speech. However, we were also interested in whether the latency of peak feature encoding varied by acoustic feature, as this would further support the independence of their contributions. We perform this analysis using the encoding weights for unprocessed English only.

For each feature, we used the MATLAB function `islocalmax` to identify the latency of peak encoding weights on a per-participant basis. The acoustic features (Main manuscript, Figure 3, Panel A) are characterised by an early positive peak, ranging from Spectral Flux (Median latency $= 90.00$ ms, 95% CI $[80.00, 90.00]$, IQR $= 20.00$) and Envelope Derivative (Median latency $= 90.00$ ms, 95% CI $[80.00, 140.00]$, IQR $= 130.00$), to Spectral Variation (Median latency $= 105.00$ ms, 95% CI $[100.00, 115.00]$, IQR $= 20.00$), and finally, Envelope (Median latency $= 200.00$ ms (95% CI $[170.00, 240.00]$, IQR $= 230.00$). Spectral Flux and Envelope Derivative also show a secondary positive peak occurring from approximately 160 to 220 ms post-stimulus. In general, the positive acoustic feature weights are strongly left-lateralised, with a corresponding negativity in right sensor space at similar time points.

In comparison to the acoustic features, the linguistic features (Main manuscript, Figure 3, Panel B) evoke more diffuse weights over time. The first peak is for Phoneme Onsets (Median latency $= 145.00$ ms, 95% CI $[100.00, 210.00]$, IQR $= 180.00$), followed by Syllable Onsets (Median latency $= 240.00$ ms, 95% CI $[93.00, 375.00]$, IQR $= 180.00$), and Word Onsets (Median latency $= 425.00$ ms, 95% CI $[370.00, 440.00]$, IQR $= 120.00$).

### 4 Linear Mixed Effect Modelling: Temporal Clusters

| Cluster | | | Time (ms) | | | | $T$-Statistics | | |
|---|---|---|---|---|---|---|---|---|---|
| Feature | ROI | Effect | Start | Max | End | Length | Sum | Max | $p_{cluster}$ |
| EnvDeriv | CF | Unprocessed / Vocoded | 140 | 150 | 270 | 130 | 40.47 | 3.77 | 0.0024 |
| EnvDeriv | CF | Unprocessed / Vocoded | -20 | 20 | 60 | 80 | 39.57 | 5.73 | 0.0024 |
| EnvDeriv | CF | Unprocessed / Voc. + Blurring | 160 | 190 | 220 | 60 | 24.16 | 4.42 | 0.0060 |
| EnvDeriv | CF | Dutch x Vocoded | 590 | 640 | 650 | 60 | 33.46 | 7.4 | 0.0050 |
| EnvDeriv | CF | Dutch x Voc. + Blurring | 160 | 210 | 230 | 70 | 36.46 | 6.07 | 0.0002 |
| EnvDeriv | CP | English / Dutch | 440 | 600 | 620 | 180 | 54.99 | 3.89 | < 0.0001 |
| EnvDeriv | CP | Unprocessed / Vocoded | -20 | 20 | 60 | 80 | 35.17 | 4.8 | 0.0004 |
| EnvDeriv | CP | Unprocessed / Voc. + Blurring | -10 | 10 | 50 | 60 | 20.41 | 3.78 | 0.0008 |
| EnvDeriv | CP | Dutch x Vocoded | 570 | 650 | 650 | 80 | 31.81 | 5.81 | < 0.0001 |
| EnvDeriv | CP | Dutch x Voc. + Blurring | 170 | 210 | 220 | 50 | 17.72 | 3.46 | 0.0008 |
| EnvDeriv | LF | English / Dutch | 440 | 600 | 650 | 210 | 77.86 | 4.96 | 0.0008 |
| EnvDeriv | LF | Unprocessed / Vocoded | -30 | 20 | 50 | 80 | 49.34 | 6.72 | 0.0008 |
| EnvDeriv | LF | Unprocessed / Voc. + Blurring | -20 | 20 | 60 | 80 | 41.25 | 6.62 | 0.0002 |
| EnvDeriv | LF | Dutch x Vocoded | 10 | 60 | 170 | 160 | 67.45 | 6.11 | < 0.0001 |
| EnvDeriv | LF | Dutch x Vocoded | 570 | 650 | 650 | 80 | 38.18 | 7.9 | 0.0040 |
| EnvDeriv | LP | English / Dutch | 430 | 600 | 650 | 220 | 82.13 | 5.69 | 0.0016 |
| EnvDeriv | LP | Unprocessed / Vocoded | 140 | 150 | 270 | 130 | 41.98 | 4.17 | 0.0040 |
| EnvDeriv | LP | Unprocessed / Vocoded | -20 | 20 | 60 | 80 | 41.79 | 6.11 | 0.0042 |
| EnvDeriv | LP | Unprocessed / Voc. + Blurring | 0 | 20 | 70 | 70 | 30.45 | 4.96 | 0.0008 |
| EnvDeriv | LP | Dutch x Vocoded | 10 | 50 | 170 | 160 | 63 | 5.28 | < 0.0001 |
| EnvDeriv | LP | Dutch x Voc. + Blurring | 150 | 200 | 230 | 80 | 37.9 | 5.98 | 0.0014 |
| EnvDeriv | RF | Unprocessed / Vocoded | 30 | 120 | 150 | 120 | 41.51 | 4.46 | < 0.0001 |
| EnvDeriv | RF | Unprocessed / Voc. + Blurring | 90 | 110 | 190 | 100 | 32.89 | 3.54 | < 0.0001 |
| EnvDeriv | RP | Unprocessed / Vocoded | 40 | 130 | 160 | 120 | 44.38 | 4.81 | < 0.0001 |
| EnvDeriv | RP | Unprocessed / Voc. + Blurring | 90 | 160 | 200 | 110 | 45.34 | 4.73 | < 0.0001 |
| EnvDeriv | RP | Dutch x Vocoded | 130 | 160 | 190 | 60 | 24.63 | 4.27 | < 0.0001 |
| EnvDeriv | RP | Dutch x Voc. + Blurring | 170 | 190 | 220 | 50 | 19.42 | 4.17 | 0.0002 |
| Envelope | CF | English / Dutch | -100 | -70 | -30 | 70 | 41.76 | 6.93 | < 0.0001 |
| Envelope | CF | Unprocessed / Vocoded | -150 | -60 | 40 | 190 | 69.72 | 5.26 | 0.0004 |
| Envelope | CF | Unprocessed / Vocoded | 210 | 230 | 350 | 140 | 76.07 | 7.11 | < 0.0001 |
| Envelope | CF | Unprocessed / Voc. + Blurring | 220 | 290 | 370 | 150 | 95.23 | 9.5 | < 0.0001 |
| Envelope | CF | Dutch x Vocoded | -110 | -80 | -40 | 70 | 32.96 | 5.16 | < 0.0001 |
| Envelope | CF | Dutch x Voc. + Blurring | -100 | -80 | -40 | 60 | 27.95 | 4.87 | 0.0002 |
| Envelope | CP | English / Dutch | -100 | -70 | -30 | 70 | 30.43 | 4.79 | < 0.0001 |
| Envelope | CP | Unprocessed / Vocoded | -120 | -60 | 30 | 150 | 50.08 | 4.22 | < 0.0001 |
| Envelope | CP | Unprocessed / Vocoded | 210 | 230 | 330 | 120 | 48.75 | 4.68 | 0.0002 |
| Envelope | CP | Unprocessed / Voc. + Blurring | 230 | 290 | 370 | 140 | 76.59 | 7.16 | < 0.0001 |
| Envelope | CP | Dutch x Vocoded | -110 | -90 | -40 | 70 | 24.86 | 3.8 | < 0.0001 |
| Envelope | CP | Dutch x Voc. + Blurring | -100 | -80 | -40 | 60 | 20.88 | 3.53 | < 0.0001 |
| Envelope | LF | English / Dutch | 90 | 130 | 160 | 70 | 31 | 5.11 | 0.0070 |
| Envelope | LF | English / Dutch | -110 | -60 | -10 | 100 | 57.33 | 6.81 | < 0.0001 |
| Envelope | LF | Unprocessed / Vocoded | -150 | -10 | 50 | 200 | 91.31 | 6.6 | 0.0002 |
| Envelope | LF | Unprocessed / Vocoded | 190 | 240 | 360 | 170 | 112.55 | 10.09 | < 0.0001 |
| Envelope | LF | Unprocessed / Voc. + Blurring | 210 | 290 | 380 | 170 | 137.13 | 12.45 | 0.0002 |
| Envelope | LF | Dutch x Vocoded | -110 | -90 | -40 | 70 | 26.37 | 4.15 | 0.0016 |
| Envelope | LF | Dutch x Voc. + Blurring | -100 | -70 | -30 | 70 | 35.08 | 5.21 | < 0.0001 |
| Envelope | LP | English / Dutch | -110 | -70 | -30 | 80 | 47.08 | 6.84 | < 0.0001 |
| Envelope | LP | Unprocessed / Vocoded | -150 | -50 | 30 | 180 | 71.88 | 5.27 | 0.0006 |
| Envelope | LP | Unprocessed / Vocoded | 200 | 240 | 370 | 170 | 104.86 | 9.15 | < 0.0001 |
| Envelope | LP | Unprocessed / Voc. + Blurring | 220 | 290 | 380 | 160 | 121.25 | 11.29 | < 0.0001 |
| Envelope | LP | Dutch x Vocoded | -110 | -90 | -30 | 80 | 37.63 | 5.45 | 0.0002 |
| Envelope | LP | Dutch x Voc. + Blurring | -110 | -90 | -40 | 70 | 30.18 | 4.74 | < 0.0001 |
| Envelope | RF | English / Dutch | 220 | 340 | 400 | 180 | 55.79 | 3.86 | < 0.0001 |
| Envelope | RF | Unprocessed / Voc. + Blurring | -10 | 20 | 40 | 50 | 17.96 | 3.44 | < 0.0001 |
| Envelope | RF | Dutch x Voc. + Blurring | 330 | 360 | 380 | 50 | 17.09 | 3.36 | 0.0056 |
| Envelope | RP | English / Dutch | 300 | 340 | 360 | 60 | 20.08 | 3.5 | 0.0036 |
| Envelope | RP | Unprocessed / Vocoded | 120 | 140 | 170 | 50 | 18.14 | 3.7 | < 0.0001 |
| PhonemeOnsets | CF | English / Dutch | 460 | 630 | 650 | 190 | 80.37 | 6.03 | < 0.0001 |

Continued on next page

**Table 1 – continued from previous page**

| Feature | ROI | Effect | Start | Max | End | Length | Sum | Max | $p$ |
|---|---|---|---|---|---|---|---|---|---|
| PhonemeOnsets | CF | Unprocessed / Vocoded | 70 | 210 | 360 | 290 | 146.4 | 6.68 | < 0.0001 |
| PhonemeOnsets | CF | Unprocessed / Voc. + Blurring | 420 | 610 | 650 | 230 | 98.07 | 6.6 | < 0.0001 |
| PhonemeOnsets | CF | Unprocessed / Voc. + Blurring | 60 | 210 | 330 | 270 | 139.7 | 8.18 | < 0.0001 |
| PhonemeOnsets | CF | Dutch x Vocoded | 0 | 110 | 230 | 230 | 79.13 | 3.95 | < 0.0001 |
| PhonemeOnsets | CF | Dutch x Voc. + Blurring | 250 | 500 | 540 | 290 | 90.29 | 4.17 | < 0.0001 |
| PhonemeOnsets | CP | English / Dutch | 460 | 630 | 650 | 190 | 82.12 | 5.59 | < 0.0001 |
| PhonemeOnsets | CP | Unprocessed / Vocoded | 100 | 220 | 350 | 250 | 80.28 | 3.76 | < 0.0001 |
| PhonemeOnsets | CP | Unprocessed / Vocoded | 550 | 620 | 650 | 100 | 45.01 | 5.12 | < 0.0001 |
| PhonemeOnsets | CP | Unprocessed / Voc. + Blurring | 140 | 210 | 320 | 180 | 82.48 | 5.89 | < 0.0001 |
| PhonemeOnsets | CP | Unprocessed / Voc. + Blurring | 430 | 610 | 650 | 220 | 91.55 | 5.94 | < 0.0001 |
| PhonemeOnsets | CP | Dutch x Vocoded | 290 | 320 | 360 | 70 | 22.13 | 3.38 | 0.0028 |
| PhonemeOnsets | CP | Dutch x Vocoded | 590 | 630 | 650 | 60 | 24.46 | 4.08 | 0.0006 |
| PhonemeOnsets | CP | Dutch x Voc. + Blurring | 430 | 610 | 650 | 220 | 80.92 | 4.74 | < 0.0001 |
| PhonemeOnsets | LF | English / Dutch | 580 | 630 | 650 | 70 | 31.41 | 4.89 | 0.0028 |
| PhonemeOnsets | LF | English / Dutch | 270 | 320 | 370 | 100 | 40.28 | 4.98 | < 0.0001 |
| PhonemeOnsets | LF | Unprocessed / Vocoded | 90 | 210 | 370 | 280 | 173.51 | 8.45 | < 0.0001 |
| PhonemeOnsets | LF | Unprocessed / Voc. + Blurring | 150 | 220 | 340 | 190 | 140.48 | 10 | 0.0010 |
| PhonemeOnsets | LF | Dutch x Vocoded | 70 | 200 | 230 | 160 | 50.44 | 3.55 | < 0.0001 |
| PhonemeOnsets | LF | Dutch x Vocoded | 280 | 320 | 370 | 90 | 35.93 | 4.65 | 0.0016 |
| PhonemeOnsets | LF | Dutch x Voc. + Blurring | -10 | 70 | 130 | 140 | 53.58 | 4.62 | 0.0002 |
| PhonemeOnsets | LF | Dutch x Voc. + Blurring | 410 | 500 | 650 | 240 | 91.58 | 5.06 | < 0.0001 |
| PhonemeOnsets | LP | English / Dutch | 0 | 190 | 230 | 230 | 63.86 | 3.63 | 0.0004 |
| PhonemeOnsets | LP | English / Dutch | 440 | 630 | 650 | 210 | 89.57 | 6.03 | < 0.0001 |
| PhonemeOnsets | LP | Unprocessed / Vocoded | 490 | 620 | 650 | 160 | 68.65 | 6.13 | 0.0026 |
| PhonemeOnsets | LP | Unprocessed / Vocoded | 60 | 210 | 360 | 300 | 159.02 | 7.45 | < 0.0001 |
| PhonemeOnsets | LP | Unprocessed / Voc. + Blurring | 410 | 610 | 650 | 240 | 135.81 | 7.87 | < 0.0001 |
| PhonemeOnsets | LP | Unprocessed / Voc. + Blurring | 70 | 210 | 330 | 260 | 143.64 | 8.74 | < 0.0001 |
| PhonemeOnsets | LP | Dutch x Vocoded | 0 | 110 | 230 | 230 | 82.62 | 4.29 | < 0.0001 |
| PhonemeOnsets | LP | Dutch x Voc. + Blurring | 310 | 500 | 650 | 340 | 131.5 | 5.6 | < 0.0001 |
| PhonemeOnsets | RF | English / Dutch | -90 | 110 | 230 | 320 | 139.14 | 6.64 | < 0.0001 |
| PhonemeOnsets | RF | English / Dutch | 460 | 630 | 650 | 190 | 104.63 | 7.19 | 0.0004 |
| PhonemeOnsets | RF | Unprocessed / Vocoded | 570 | 620 | 650 | 80 | 31.79 | 4.14 | < 0.0001 |
| PhonemeOnsets | RF | Unprocessed / Voc. + Blurring | 540 | 610 | 650 | 110 | 45.57 | 4.93 | 0.0038 |
| PhonemeOnsets | RF | Unprocessed / Voc. + Blurring | 40 | 100 | 200 | 160 | 63.27 | 4.99 | < 0.0001 |
| PhonemeOnsets | RF | Dutch x Vocoded | 580 | 630 | 650 | 70 | 27.63 | 4.2 | 0.0006 |
| PhonemeOnsets | RF | Dutch x Voc. + Blurring | 30 | 90 | 120 | 90 | 30.21 | 3.78 | 0.0058 |
| PhonemeOnsets | RF | Dutch x Voc. + Blurring | 530 | 620 | 650 | 120 | 54.78 | 5.72 | < 0.0001 |
| PhonemeOnsets | RP | English / Dutch | 470 | 630 | 650 | 180 | 81.56 | 5.79 | < 0.0001 |
| PhonemeOnsets | RP | English / Dutch | -40 | 90 | 200 | 240 | 94.53 | 6.05 | < 0.0001 |
| PhonemeOnsets | RP | Unprocessed / Vocoded | 560 | 620 | 650 | 90 | 34.37 | 4.1 | < 0.0001 |
| PhonemeOnsets | RP | Unprocessed / Voc. + Blurring | 50 | 120 | 240 | 190 | 85.76 | 5.4 | < 0.0001 |
| PhonemeOnsets | RP | Unprocessed / Voc. + Blurring | 350 | 610 | 650 | 300 | 119.39 | 6.23 | < 0.0001 |
| PhonemeOnsets | RP | Dutch x Voc. + Blurring | 240 | 350 | 410 | 170 | 58.35 | 3.78 | < 0.0001 |
| SpectralFlux | CF | Unprocessed / Vocoded | 330 | 430 | 450 | 120 | 60.76 | 7.69 | < 0.0001 |
| SpectralFlux | CF | Unprocessed / Vocoded | 40 | 70 | 110 | 70 | 42.47 | 8.07 | 0.0046 |
| SpectralFlux | CF | Unprocessed / Voc. + Blurring | 340 | 390 | 530 | 190 | 82.34 | 7.08 | 0.0004 |
| SpectralFlux | CP | English / Dutch | 530 | 560 | 580 | 50 | 28.23 | 5.74 | 0.0072 |
| SpectralFlux | CP | Unprocessed / Vocoded | 40 | 70 | 100 | 60 | 24.23 | 4.64 | 0.0008 |
| SpectralFlux | CP | Unprocessed / Vocoded | 380 | 430 | 440 | 60 | 25.48 | 4.79 | 0.0002 |
| SpectralFlux | CP | Unprocessed / Voc. + Blurring | 350 | 400 | 440 | 90 | 33.13 | 4.79 | 0.0012 |
| SpectralFlux | LF | Unprocessed / Vocoded | 20 | 70 | 160 | 140 | 91.86 | 13.33 | 0.0002 |
| SpectralFlux | LP | Unprocessed / Vocoded | 340 | 430 | 450 | 110 | 55.99 | 7.84 | 0.0010 |
| SpectralFlux | RF | English / Dutch | -120 | -100 | -50 | 70 | 27.1 | 5.06 | 0.0020 |
| SpectralFlux | RF | Unprocessed / Voc. + Blurring | 50 | 80 | 100 | 50 | 22.06 | 4.87 | 0.0002 |
| SpectralFlux | RP | Unprocessed / Vocoded | 360 | 400 | 440 | 80 | 29.89 | 4.36 | < 0.0001 |
| SpectralFlux | RP | Unprocessed / Voc. + Blurring | 350 | 390 | 420 | 70 | 26.85 | 5.03 | 0.0002 |
| SpectralFlux | RP | Dutch x Vocoded | 380 | 400 | 430 | 50 | 18.74 | 4.03 | 0.0010 |
| SpectralVariation | CF | English / Dutch | 370 | 440 | 530 | 160 | 75.92 | 7.67 | < 0.0001 |
| SpectralVariation | CF | Unprocessed / Voc. + Blurring | 130 | 250 | 340 | 210 | 92.95 | 6.53 | 0.0010 |
| SpectralVariation | CF | Dutch x Vocoded | 420 | 450 | 540 | 120 | 56.18 | 5.45 | < 0.0001 |
| SpectralVariation | CF | Dutch x Voc. + Blurring | -30 | -10 | 110 | 140 | 48.84 | 4.43 | 0.0006 |

Continued on next page

**Table 1 – continued from previous page**

| Feature | ROI | Effect | Start | Max | End | Length | Sum | Max | *p* |
|---|---|---|---|---|---|---|---|---|---|
| SpectralVariation | CF | Dutch x Voc. + Blurring | 420 | 440 | 530 | 110 | 43.86 | 6.56 | 0.0024 |
| SpectralVariation | CP | English / Dutch | 370 | 440 | 530 | 160 | 58.56 | 5.98 | < 0.0001 |
| SpectralVariation | CP | Unprocessed / Voc. + Blurring | 140 | 250 | 280 | 140 | 51.8 | 4.29 | 0.0002 |
| SpectralVariation | CP | Dutch x Vocoded | 430 | 450 | 540 | 110 | 41.66 | 5.07 | < 0.0001 |
| SpectralVariation | CP | Dutch x Voc. + Blurring | 420 | 440 | 530 | 110 | 38.59 | 5.19 | < 0.0001 |
| SpectralVariation | LF | English / Dutch | 360 | 440 | 530 | 170 | 77.96 | 7.75 | < 0.0001 |
| SpectralVariation | LF | Unprocessed / Voc. + Blurring | 170 | 260 | 350 | 180 | 125.81 | 8.8 | < 0.0001 |
| SpectralVariation | LF | Dutch x Vocoded | 420 | 450 | 530 | 110 | 49.89 | 6.32 | < 0.0001 |
| SpectralVariation | LF | Dutch x Voc. + Blurring | -30 | -10 | 100 | 130 | 57.04 | 6.08 | 0.0004 |
| SpectralVariation | LF | Dutch x Voc. + Blurring | 420 | 440 | 520 | 100 | 42.86 | 6.68 | 0.0072 |
| SpectralVariation | LP | English / Dutch | 370 | 440 | 530 | 160 | 79.24 | 7.93 | 0.0002 |
| SpectralVariation | LP | Unprocessed / Voc. + Blurring | 130 | 210 | 340 | 210 | 111.16 | 7.63 | 0.0004 |
| SpectralVariation | LP | Dutch x Vocoded | 420 | 450 | 540 | 120 | 54.86 | 5.82 | < 0.0001 |
| SpectralVariation | LP | Dutch x Voc. + Blurring | -30 | -10 | 110 | 140 | 61.91 | 5.32 | 0.0002 |
| SpectralVariation | RF | English / Dutch | 210 | 250 | 270 | 60 | 21.87 | 4.14 | 0.0008 |
| SpectralVariation | RF | Unprocessed / Vocoded | 240 | 270 | 290 | 50 | 22.25 | 4.83 | 0.0006 |
| SpectralVariation | RF | Unprocessed / Voc. + Blurring | 210 | 280 | 350 | 140 | 51.37 | 4.74 | < 0.0001 |
| SpectralVariation | RP | English / Dutch | 170 | 240 | 260 | 90 | 30.42 | 3.7 | 0.0006 |
| SpectralVariation | RP | Unprocessed / Vocoded | 230 | 260 | 290 | 60 | 25.92 | 4.31 | 0.0010 |
| SpectralVariation | RP | Dutch x Vocoded | 160 | 250 | 280 | 120 | 39.36 | 4.1 | < 0.0001 |
| SpectralVariation | RP | Dutch x Voc. + Blurring | 180 | 200 | 250 | 70 | 22.81 | 3.48 | 0.0052 |
| SyllableOnsets | CF | English / Dutch | 400 | 450 | 540 | 140 | 46.94 | 3.93 | < 0.0001 |
| SyllableOnsets | CF | Unprocessed / Vocoded | -150 | 260 | 320 | 470 | 134.19 | 3.51 | < 0.0001 |
| SyllableOnsets | CF | Unprocessed / Voc. + Blurring | -150 | -50 | 40 | 190 | 61.25 | 3.69 | < 0.0001 |
| SyllableOnsets | CF | Unprocessed / Voc. + Blurring | 320 | 420 | 520 | 200 | 68.66 | 4.09 | < 0.0001 |
| SyllableOnsets | CF | Dutch x Vocoded | -60 | 250 | 320 | 380 | 116.78 | 3.37 | < 0.0001 |
| SyllableOnsets | CF | Dutch x Voc. + Blurring | 280 | 400 | 510 | 230 | 81.53 | 4.43 | < 0.0001 |
| SyllableOnsets | LF | Unprocessed / Vocoded | 60 | 260 | 330 | 270 | 93.49 | 4.19 | < 0.0001 |
| SyllableOnsets | LF | Unprocessed / Voc. + Blurring | 320 | 430 | 570 | 250 | 98.63 | 4.97 | < 0.0001 |
| SyllableOnsets | LF | Dutch x Voc. + Blurring | 250 | 400 | 520 | 270 | 102.13 | 4.87 | < 0.0001 |
| SyllableOnsets | LP | English / Dutch | 400 | 450 | 520 | 120 | 43.39 | 4.19 | < 0.0001 |
| SyllableOnsets | LP | Unprocessed / Vocoded | -150 | 250 | 330 | 480 | 175.38 | 4.5 | 0.0006 |
| SyllableOnsets | LP | Unprocessed / Voc. + Blurring | -150 | -50 | 220 | 370 | 107.26 | 3.9 | 0.0004 |
| SyllableOnsets | LP | Unprocessed / Voc. + Blurring | 320 | 440 | 560 | 240 | 102.87 | 5.47 | 0.0004 |
| SyllableOnsets | LP | Dutch x Vocoded | -120 | 240 | 320 | 440 | 132.95 | 3.51 | < 0.0001 |
| SyllableOnsets | LP | Dutch x Voc. + Blurring | 290 | 400 | 520 | 230 | 87.8 | 4.74 | < 0.0001 |
| SyllableOnsets | RP | Dutch x Voc. + Blurring | 550 | 610 | 650 | 100 | 34.42 | 3.76 | < 0.0001 |
| WordOnsets | CF | English / Dutch | 250 | 420 | 650 | 400 | 262.9 | 10.25 | < 0.0001 |
| WordOnsets | CF | Unprocessed / Vocoded | 280 | 420 | 460 | 180 | 62.25 | 3.75 | < 0.0001 |
| WordOnsets | CF | Unprocessed / Vocoded | -150 | -80 | -10 | 140 | 69.27 | 5.51 | < 0.0001 |
| WordOnsets | CF | Unprocessed / Voc. + Blurring | 260 | 420 | 650 | 390 | 297.21 | 11.63 | < 0.0001 |
| WordOnsets | CF | Dutch x Vocoded | -150 | -90 | -50 | 100 | 35.39 | 3.91 | < 0.0001 |
| WordOnsets | CF | Dutch x Voc. + Blurring | 270 | 420 | 650 | 380 | 158.62 | 5.8 | < 0.0001 |
| WordOnsets | CP | English / Dutch | 270 | 420 | 650 | 380 | 213.57 | 7.86 | < 0.0001 |
| WordOnsets | CP | Unprocessed / Vocoded | 270 | 320 | 460 | 190 | 59.61 | 3.45 | < 0.0001 |
| WordOnsets | CP | Unprocessed / Voc. + Blurring | 270 | 430 | 650 | 380 | 243.11 | 8.99 | < 0.0001 |
| WordOnsets | CP | Dutch x Voc. + Blurring | 280 | 420 | 650 | 370 | 151.54 | 5.19 | < 0.0001 |
| WordOnsets | LF | English / Dutch | 180 | 420 | 650 | 470 | 197.69 | 6.84 | < 0.0001 |
| WordOnsets | LF | Unprocessed / Vocoded | 280 | 430 | 470 | 190 | 61.08 | 3.65 | < 0.0001 |
| WordOnsets | LF | Unprocessed / Voc. + Blurring | 240 | 430 | 650 | 410 | 257.74 | 8.72 | < 0.0001 |
| WordOnsets | LF | Dutch x Voc. + Blurring | 400 | 650 | 650 | 250 | 83.8 | 4.18 | < 0.0001 |
| WordOnsets | LP | English / Dutch | 250 | 420 | 650 | 400 | 267.69 | 10.4 | 0.0002 |
| WordOnsets | LP | Unprocessed / Vocoded | 220 | 320 | 460 | 240 | 84.12 | 4.51 | < 0.0001 |
| WordOnsets | LP | Unprocessed / Vocoded | -150 | -80 | -10 | 140 | 67.14 | 5.37 | < 0.0001 |
| WordOnsets | LP | Unprocessed / Voc. + Blurring | 260 | 430 | 650 | 390 | 329.43 | 12.53 | < 0.0001 |
| WordOnsets | LP | Dutch x Vocoded | -150 | -90 | -40 | 110 | 41.55 | 4.29 | < 0.0001 |
| WordOnsets | LP | Dutch x Voc. + Blurring | 260 | 420 | 650 | 390 | 164.96 | 5.92 | < 0.0001 |
| WordOnsets | RF | English / Dutch | 290 | 420 | 650 | 360 | 225.49 | 7.99 | 0.0002 |
| WordOnsets | RF | Unprocessed / Voc. + Blurring | 300 | 420 | 650 | 350 | 191.61 | 7.93 | < 0.0001 |
| WordOnsets | RF | Dutch x Voc. + Blurring | 270 | 410 | 650 | 380 | 180.95 | 6.42 | < 0.0001 |
| WordOnsets | RP | Unprocessed / Vocoded | 280 | 390 | 460 | 180 | 56.84 | 3.34 | < 0.0001 |

Continued on next page

**Table 1 – continued from previous page**

| Feature | ROI | Effect | Start | Max | End | Length | Sum | Max | $p$ |
|---|---|---|---|---|---|---|---|---|---|
| WordOnsets | RP | Unprocessed / Voc. + Blurring | 270 | 420 | 650 | 380 | 236.91 | 9.37 | < 0.0001 |
| WordOnsets | RP | Dutch x Vocoded | 250 | 310 | 430 | 180 | 51.5 | 3.5 | < 0.0001 |
| WordOnsets | RP | Dutch x Voc. + Blurring | 260 | 410 | 650 | 390 | 191.26 | 6.69 | < 0.0001 |

Table 1: Clusters identified using mass permutation tests with 5000 iterations. Abbreviations: CF Central Frontal; LF Left Frontal; RF Right Frontal; CP Central Posterior; LP Left Posterior; RP Right Posterior. The threshold of significance for multiple comparisons over six regions of interest is set at $p_{cluster} \leq 0.008$ .

### 5 Linear Mixed Effect Modelling: Results Visualised by Sensor Region of Interest

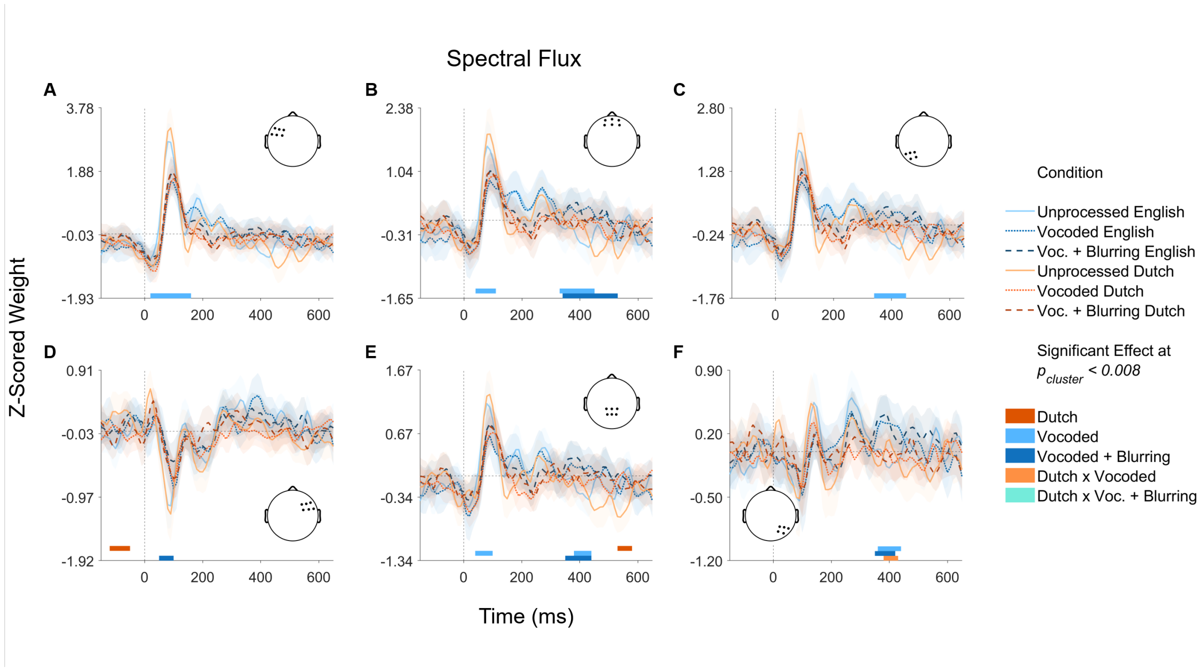


Figure 7: Z-scored encoding model weights across experimental conditions for Spectral Flux with cluster-level results for linear mixed effect models. The model reference level is unprocessed English. Coloured bars indicate time points where a significant effect was found according to cluster-based permutation testing (5,000 iterations). Traces and shaded regions indicate the group mean (averaged over sensors) and 95% confidence intervals of the mean.

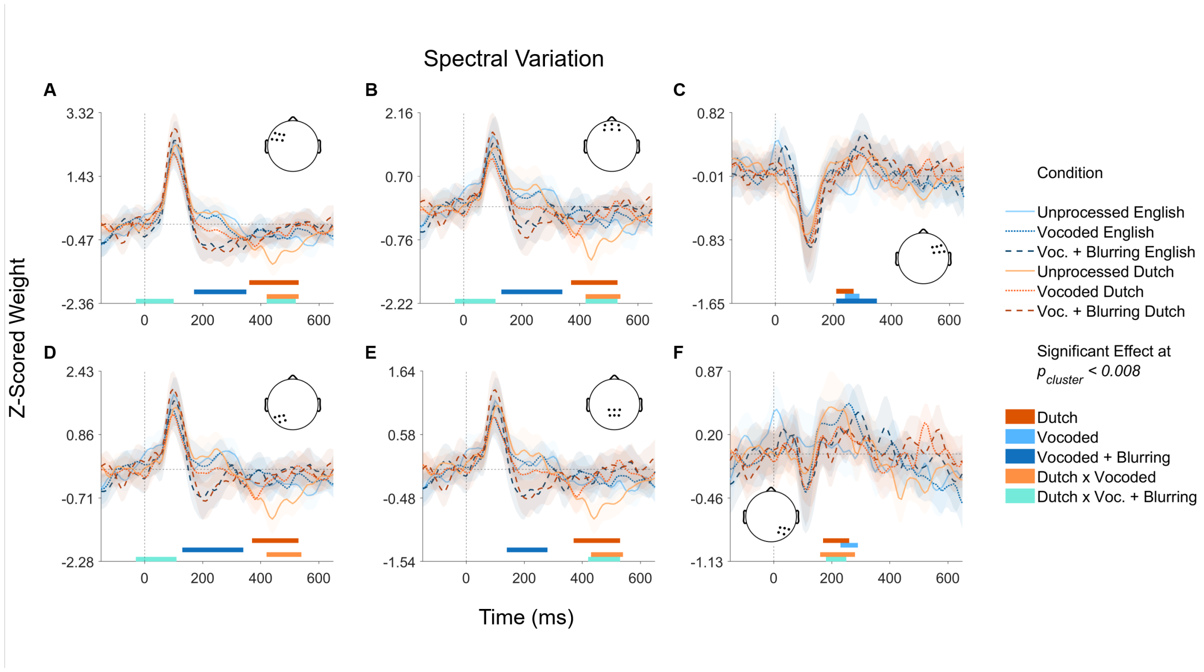


Figure 8: Z-scored encoding model weights across experimental conditions for Spectral Variation with cluster-level results for linear mixed effect models. The model reference level is unprocessed English. Coloured bars indicate time points where a significant effect was found according to cluster-based permutation testing (5,000 iterations). Traces and shaded regions indicate the group mean (averaged over sensors) and 95% confidence intervals of the mean.

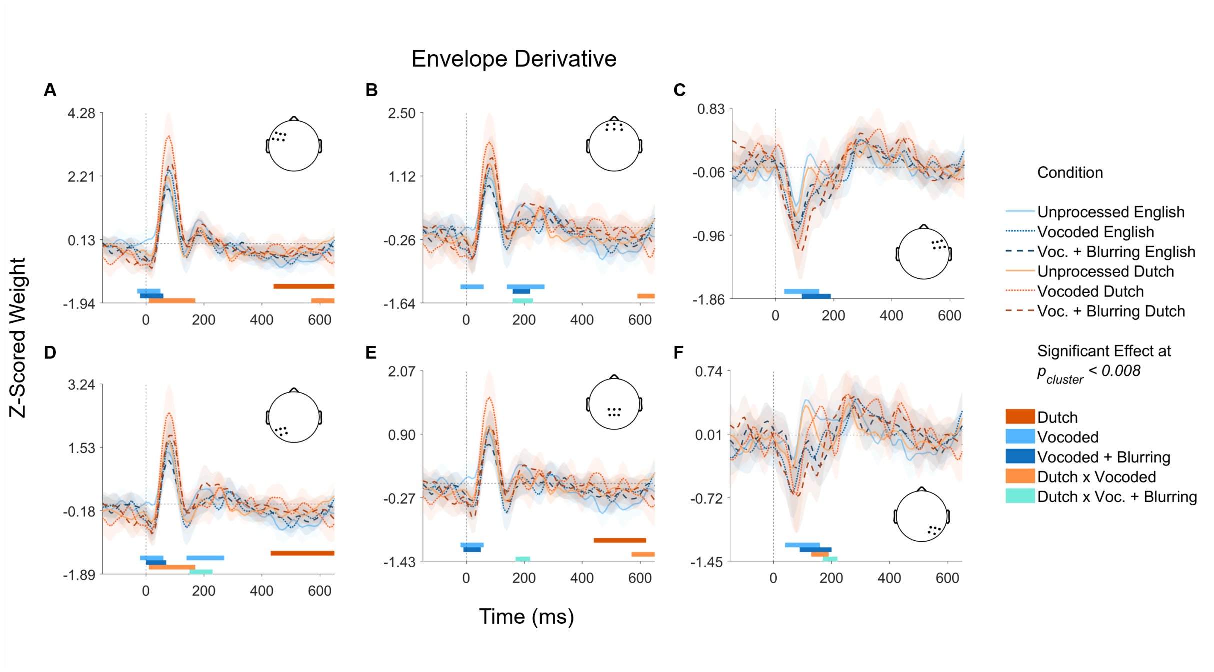


Figure 9: Z-scored encoding model weights across experimental conditions for Envelope Derivative with cluster-level results for linear mixed effect models. The model reference level is unprocessed English. Coloured bars indicate time points where a significant effect was found according to cluster-based permutation testing (5,000 iterations). Traces and shaded regions indicate the group mean (averaged over sensors) and 95% confidence intervals of the mean.

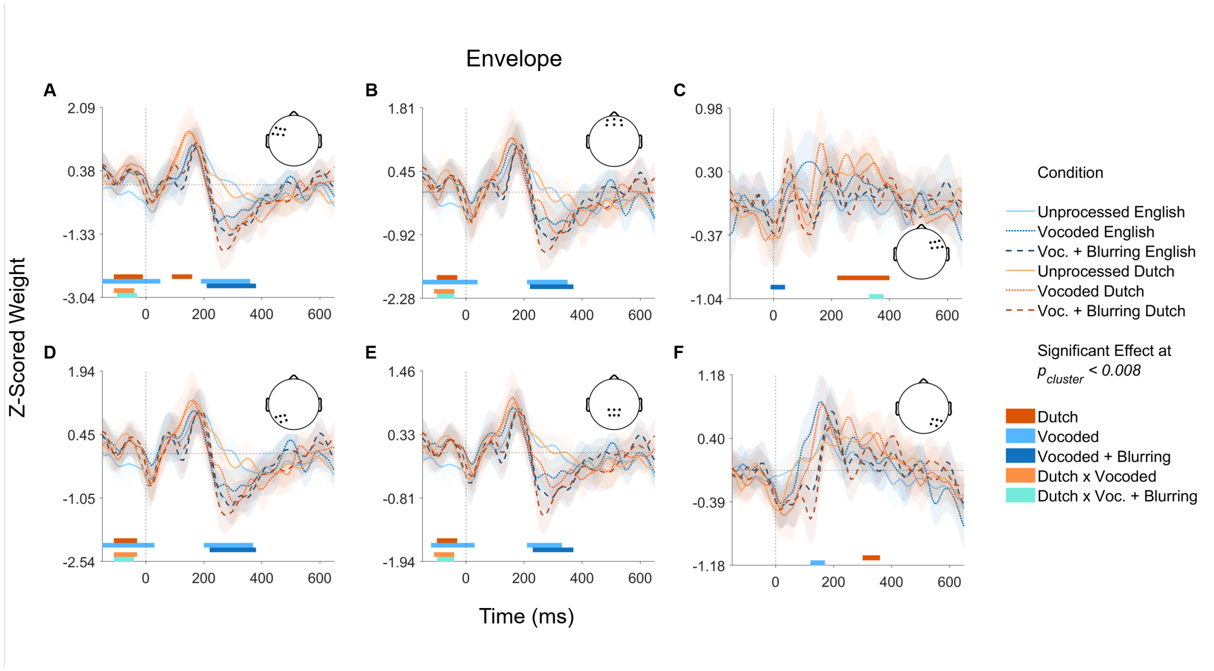


Figure 10: Z-scored encoding model weights across experimental conditions for Envelope with cluster-level results for linear mixed effect models. The model reference level is unprocessed English. Coloured bars indicate time points where a significant effect was found according to cluster-based permutation testing (5,000 iterations). Traces and shaded regions indicate the group mean (averaged over sensors) and 95% confidence intervals of the mean.

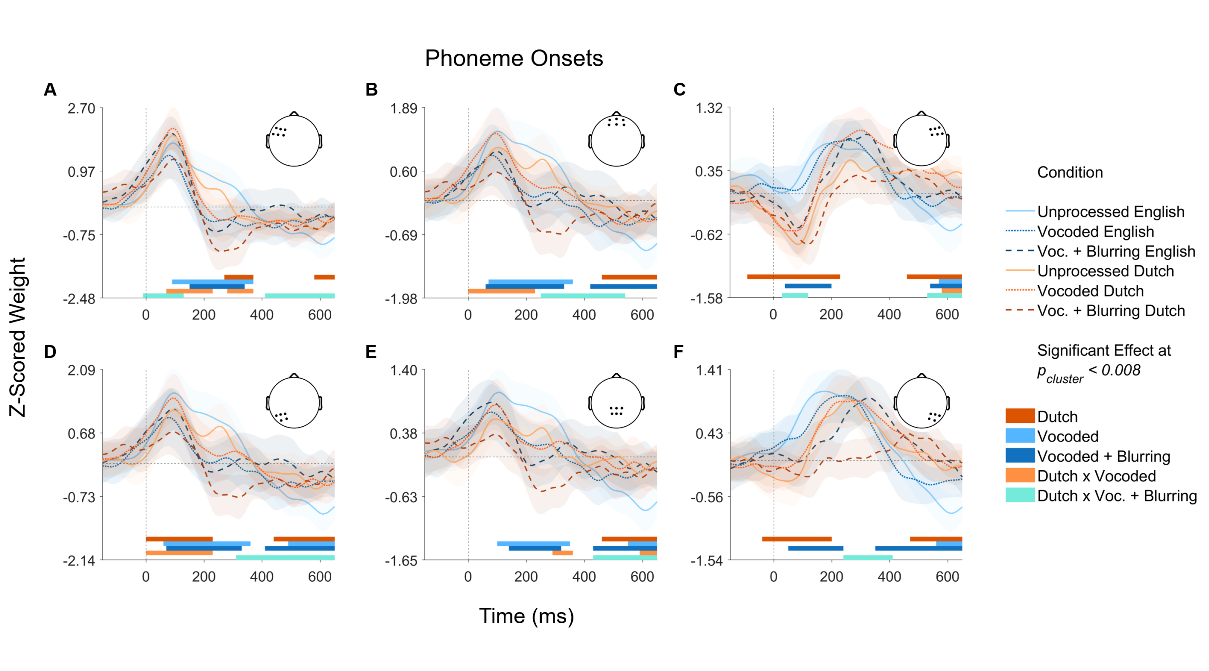


Figure 11: Z-scored encoding model weights across experimental conditions for Phoneme Onsets with cluster-level results for linear mixed effect models. The model reference level is unprocessed English. Coloured bars indicate time points where a significant effect was found according to cluster-based permutation testing (5,000 iterations). Traces and shaded regions indicate the group mean (averaged over sensors) and 95% confidence intervals of the mean.

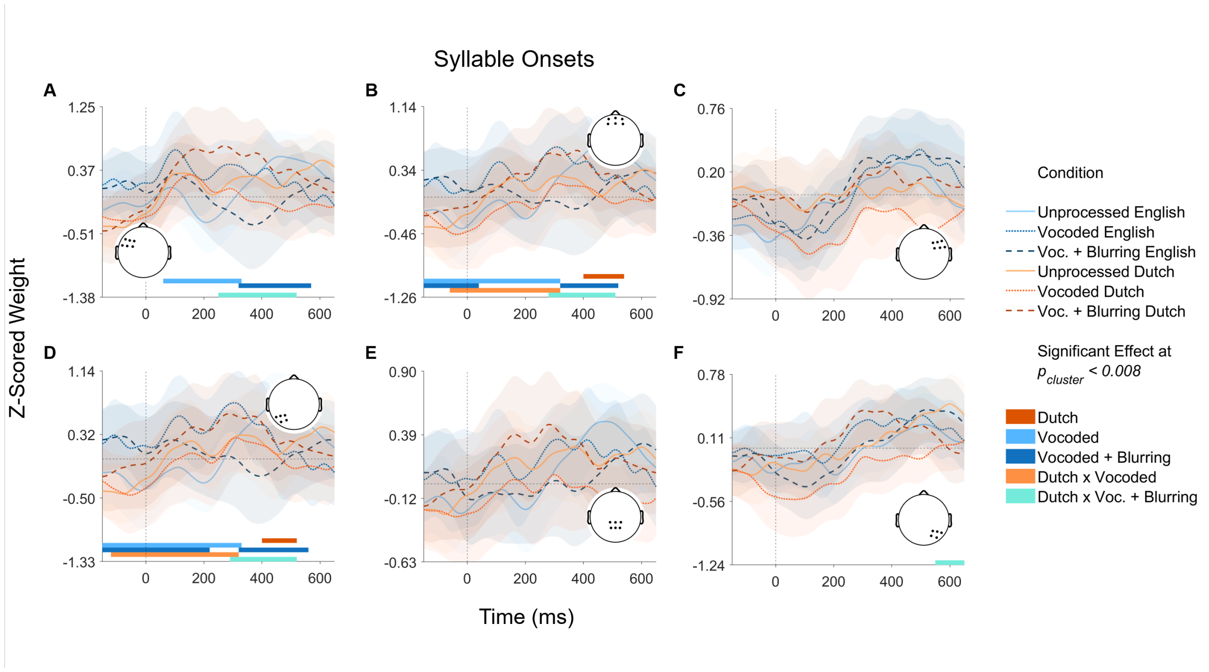


Figure 12: Z-scored encoding model weights across experimental conditions for Syllable Onsets with cluster-level results for linear mixed effect models. The model reference level is unprocessed English. Coloured bars indicate time points where a significant effect was found according to cluster-based permutation testing (5,000 iterations). Traces and shaded regions indicate the group mean (averaged over sensors) and 95% confidence intervals of the mean.

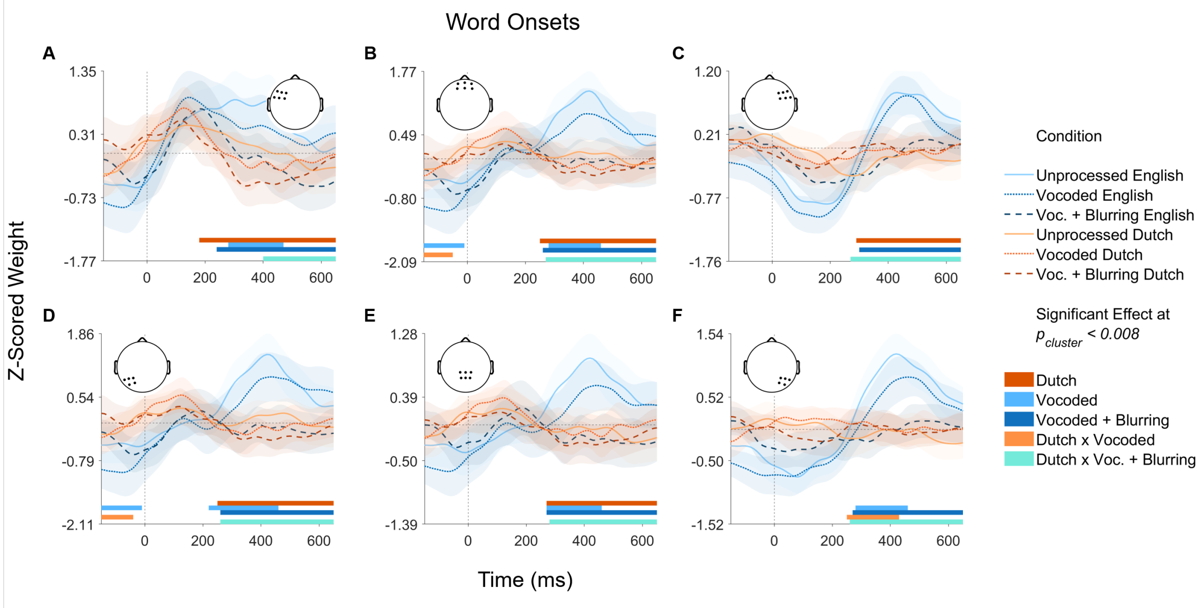


Figure 13: Z-scored encoding model weights across experimental conditions for Word Onsets with cluster-level results for linear mixed effect models. The model reference level is unprocessed English. Coloured bars indicate time points where a significant effect was found according to cluster-based permutation testing (5,000 iterations). Traces and shaded regions indicate the group mean (averaged over sensors) and 95% confidence intervals of the mean.

### 6 Linear Mixed Effect Modelling: Reported Model Details

#### 6.1 Spectral Variation - LF - $-10$ ms

Table 2: Model formula: $Z \sim$ Language $\times$ SpectralDegradation + (1 | Electrode) + (1 | Participant). Standardized parameters were obtained by fitting the model on a standardized version of the dataset. 95% Confidence Intervals (CIs) and p-values were computed using a Wald t-distribution approximation.

| | | 95% CI | | | | | | 95% CI | |
|---|---|---|---|---|---|---|---|---|---|
| Parameter | Estimate | Lower | Upper | $t$ | df | $p$ | Std Est | Lower | Upper |
| (Intercept) | 0.19 | -0.02 | 0.40 | 1.80 | 84.18 | 0.075 | 0.16 | -0.01 | 0.34 |
| Language | -0.31 | -0.51 | -0.11 | -3.01 | 1320.00 | 0.003 | -0.26 | -0.43 | -0.09 |
| Vocoded | -0.10 | -0.30 | 0.10 | -0.96 | 1320.00 | 0.336 | -0.08 | -0.25 | 0.09 |
| Vocoded + Blurring | -0.55 | -0.75 | -0.35 | -5.37 | 1320.00 | $< 0.001$ | -0.46 | -0.63 | -0.29 |
| Language×Vocoded | 0.18 | -0.11 | 0.46 | 1.22 | 1320.00 | 0.222 | 0.15 | -0.09 | 0.39 |
| Language×Vocoded + Blurring | 0.88 | 0.60 | 1.16 | 6.08 | 1320.00 | $< 0.001$ | 0.74 | 0.50 | 0.98 |

| Random Effects | Variance | SD | ICC | Model Metrics | |
|---|---|---|---|---|---|
| Within-Group Variance | 1.19 | (1.09) | | AIC | 4229.0 |
| Between-Group Variance | | | | BIC | 4275.9 |
| Random Intercept (Participant) | 0.2 | (0.45) | 0.38 | $R^2$ (Conditional) | 0.17 |
| Random Intercept (Electrode) | 0 | (0.06) | 0.05 | $R^2$ (Marginal) | 0.03 |
| | | | | ICC | 0.15 |
| N Groups | | | | RMSE | 1.08 |
| Electrode | 6 | | | | |
| Participant | 38 | | | | |
| N Observations | 1368 | | | | |

##### 6.2 Envelope Derivative - LF - 60 ms

Table 3: Model formula: $Z \sim$ Language $\times$ SpectralDegradation + (1 | Electrode) + (1 | Participant). Standardized parameters were obtained by fitting the model on a standardized version of the dataset. 95% Confidence Intervals (CIs) and p-values were computed using a Wald t-distribution approximation.

| | | 95% CI | | | | | | 95% CI | |
|---|---|---|---|---|---|---|---|---|---|
| Parameter | Estimate | Lower | Upper | *t* | df | *p* | Std Est | Lower | Upper |
| (Intercept) | -0.38 | -0.58 | -0.17 | -3.71 | 100.79 | $< 0.001$ | 0.05 | -0.11 | 0.21 |
| Language | -0.82 | -1.02 | -0.61 | -7.75 | 1320.00 | $< 0.001$ | -0.66 | -0.82 | -0.49 |
| Vocoded | 0.14 | -0.07 | 0.34 | 1.29 | 1320.00 | 0.196 | 0.11 | -0.06 | 0.28 |
| Vocoded + Blurring | 0.01 | -0.20 | 0.22 | 0.09 | 1320.00 | 0.928 | 0.01 | -0.16 | 0.17 |
| Language $\times$ Vocoded | 0.77 | 0.48 | 1.07 | 5.18 | 1320.00 | $< 0.001$ | 0.62 | 0.39 | 0.86 |
| Language $\times$ Vocoded + Blurring | 1.00 | 0.70 | 1.29 | 6.68 | 1320.00 | $< 0.001$ | 0.80 | 0.56 | 1.03 |

| Random Effects | Variance | SD | ICC | Model Metrics | |
|---|---|---|---|---|---|
| Within-Group Variance | 1.27 | (1.13) | | AIC | 4306.9 |
| Between-Group Variance | | | | BIC | 4353.9 |
| Random Intercept (Participant) | 0.18 | (0.42) | 0.35 | $R^2$ (Conditional) | 0.19 |
| Random Intercept (Electrode) | 0 | (0.02) | 0.02 | $R^2$ (Marginal) | 0.08 |
| | | | | ICC | 0.12 |
| N Groups | | | | RMSE | 1.11 |
| Electrode | 6 | | | | |
| Participant | 38 | | | | |
| N Observations | 1368 | | | | |

##### 6.3 Envelope Derivative - LP - 600 ms

Table 4: Model formula: $Z \sim$ Language $\times$ SpectralDegradation + (1 | Participant). Standardized parameters were obtained by fitting the model on a standardized version of the dataset. 95% Confidence Intervals (CIs) and p-values were computed using a Wald t-distribution approximation.

| | | 95% CI | | | | | | 95% CI | |
|---|---|---|---|---|---|---|---|---|---|
| Parameter | Estimate | Lower | Upper | $t$ | df | $p$ | Std Est | Lower | Upper |
| (Intercept) | -0.52 | -0.70 | -0.34 | -5.69 | 109.52 | $< 0.001$ | -0.25 | -0.42 | -0.08 |
| Language | 0.52 | 0.34 | 0.70 | 5.69 | 1325.00 | $< 0.001$ | 0.49 | 0.32 | 0.66 |
| Vocoded | 0.27 | 0.09 | 0.45 | 2.93 | 1325.00 | 0.003 | 0.25 | 0.08 | 0.42 |
| Vocoded + Blurring | 0.22 | 0.04 | 0.40 | 2.37 | 1325.00 | 0.018 | 0.20 | 0.03 | 0.37 |
| Language×Vocoded | -0.58 | -0.83 | -0.32 | -4.46 | 1325.00 | $< 0.001$ | -0.54 | -0.78 | -0.30 |
| Language×Vocoded + Blurring | -0.37 | -0.62 | -0.12 | -2.87 | 1325.00 | 0.004 | -0.35 | -0.59 | -0.11 |

| Random Effects | Variance | SD | ICC | Model Metrics | |
|---|---|---|---|---|---|
| Within-Group Variance | 0.95 | (0.98) | | AIC | 3916.9 |
| Between-Group Variance | | | | BIC | 3958.6 |
| Random Intercept (Participant) | 0.15 | (0.39) | 0.37 | $R^2$ (Conditional) | 0.16 |
| | | | | $R^2$ (Marginal) | 0.02 |
| N Groups | | | | ICC | 0.14 |
| Participant | 38 | | | RMSE | 0.96 |
| N Observations | 1368 | | | | |

##### 6.4 Envelope - LF - 130 ms

Table 5: Model formula: $Z \sim$ Language $\times$ SpectralDegradation + (1 | Participant). Standardized parameters were obtained by fitting the model on a standardized version of the dataset. 95% Confidence Intervals (CIs) and p-values were computed using a Wald t-distribution approximation.

| | | 95% CI | | | | | | 95% CI | |
|---|---|---|---|---|---|---|---|---|---|
| Parameter | Estimate | Lower | Upper | *t* | df | *p* | Std Est | Lower | Upper |
| (Intercept) | 0.48 | 0.13 | 0.83 | 2.77 | 69.44 | 0.007 | -0.07 | -0.27 | 0.12 |
| Language | 0.71 | 0.44 | 0.99 | 5.09 | 1325.00 | $< 0.001$ | 0.40 | 0.24 | 0.55 |
| Vocoded | 0.00 | -0.27 | 0.28 | 0.01 | 1325.00 | 0.994 | 0.00 | -0.15 | 0.15 |
| Vocoded + Blurring | -0.55 | -0.82 | -0.27 | -3.92 | 1325.00 | $< 0.001$ | -0.31 | -0.46 | -0.15 |
| Language×Vocoded | 0.07 | -0.32 | 0.46 | 0.37 | 1325.00 | 0.713 | 0.04 | -0.18 | 0.26 |
| Language×Vocoded + Blurring | -0.33 | -0.72 | 0.06 | -1.67 | 1325.00 | 0.096 | -0.18 | -0.40 | 0.03 |

| Random Effects | Variance | SD | ICC | Model Metrics | |
|---|---|---|---|---|---|
| Within-Group Variance | 2.23 | (1.49) | | AIC | 5104.9 |
| Between-Group Variance | | | | BIC | 5146.7 |
| Random Intercept (Participant) | 0.77 | (0.88) | 0.51 | $R^2$ (Conditional) | 0.31 |
| | | | | $R^2$ (Marginal) | 0.07 |
| N Groups | | | | ICC | 0.26 |
| Participant | 38 | | | RMSE | 1.47 |
| N Observations | 1368 | | | | |

#### 6.5 Phoneme Onsets - LP - 110 ms

Table 6: Model formula: $Z \sim$ Language $\times$ SpectralDegradation + (1 | Participant). Standardized parameters were obtained by fitting the model on a standardized version of the dataset. 95% Confidence Intervals (CIs) and p-values were computed using a Wald t-distribution approximation.

| | | 95% CI | | | | | | 95% CI | |
|---|---|---|---|---|---|---|---|---|---|
| Parameter | Estimate | Lower | Upper | $t$ | df | $p$ | Std Est | Lower | Upper |
| (Intercept) | 1.53 | 1.18 | 1.89 | 8.67 | 75.88 | $< 0.001$ | 0.22 | 0.03 | 0.41 |
| Language | -0.39 | -0.68 | -0.09 | -2.55 | 1325.00 | 0.011 | -0.21 | -0.37 | -0.05 |
| Vocoded | -0.68 | -0.97 | -0.38 | -4.48 | 1325.00 | $< 0.001$ | -0.37 | -0.53 | -0.21 |
| Vocoded + Blurring | -0.38 | -0.68 | -0.09 | -2.55 | 1325.00 | 0.011 | -0.21 | -0.37 | -0.05 |
| Language$\times$Vocoded | 0.91 | 0.49 | 1.33 | 4.26 | 1325.00 | $< 0.001$ | 0.49 | 0.27 | 0.72 |
| Language$\times$Vocoded + Blurring | -0.10 | -0.52 | 0.32 | -0.45 | 1325.00 | 0.650 | -0.05 | -0.28 | 0.17 |

| Random Effects | Variance | SD | ICC | Model Metrics | |
|---|---|---|---|---|---|
| Within-Group Variance | 2.6 | (1.61) | | AIC | 5305.6 |
| Between-Group Variance | | | | BIC | 5347.4 |
| Random Intercept (Participant) | 0.76 | (0.87) | 0.47 | $R^2$ (Conditional) | 0.24 |
| | | | | $R^2$ (Marginal) | 0.03 |
| N Groups | | | | ICC | 0.23 |
| Participant | 38 | | | RMSE | 1.59 |
| N Observations | 1368 | | | | |

##### 6.6 Phoneme Onsets - RF - 90 ms

Table 7: Model formula: $Z \sim$ Language $\times$ SpectralDegradation + (1 | Participant). Standardized parameters were obtained by fitting the model on a standardized version of the dataset. 95% Confidence Intervals (CIs) and p-values were computed using a Wald t-distribution approximation.

| | | 95% CI | | | | | | 95% CI | |
|---|---|---|---|---|---|---|---|---|---|
| Parameter | Estimate | Lower | Upper | $t$ | df | $p$ | Std Est | Lower | Upper |
| (Intercept) | 0.26 | -0.07 | 0.59 | 1.56 | 160.87 | 0.121 | 0.29 | 0.14 | 0.45 |
| Language | -1.03 | -1.40 | -0.66 | -5.50 | 1325.00 | $< 0.001$ | -0.49 | -0.66 | -0.31 |
| Vocoded | -0.19 | -0.56 | 0.17 | -1.03 | 1325.00 | 0.304 | -0.09 | -0.26 | 0.08 |
| Vocoded + Blurring | -0.77 | -1.14 | -0.41 | -4.13 | 1325.00 | $< 0.001$ | -0.36 | -0.54 | -0.19 |
| Language$\times$Vocoded | 0.41 | -0.11 | 0.93 | 1.56 | 1325.00 | 0.119 | 0.19 | -0.05 | 0.44 |
| Language$\times$Vocoded + Blurring | 0.86 | 0.34 | 1.38 | 3.23 | 1325.00 | 0.001 | 0.40 | 0.16 | 0.65 |

| Random Effects | Variance | SD | ICC | Model Metrics | |
|---|---|---|---|---|---|
| Within-Group Variance | 4 | (2) | | AIC | 5857.1 |
| Between-Group Variance | | | | BIC | 5898.9 |
| Random Intercept (Participant) | 0.38 | (0.62) | 0.3 | $R^2$ (Conditional) | 0.12 |
| | | | | $R^2$ (Marginal) | 0.03 |
| N Groups | | | | ICC | 0.09 |
| Participant | 38 | | | RMSE | 1.97 |
| N Observations | 1368 | | | | |

##### 6.7 Syllable Onsets - LF - 400 ms

Table 8: Model formula: $Z \sim$ Language $\times$ SpectralDegradation + (1 | Participant). Standardized parameters were obtained by fitting the model on a standardized version of the dataset. 95% Confidence Intervals (CIs) and p-values were computed using a Wald t-distribution approximation.

| | | 95% CI | | | | | | 95% CI | |
|---|---|---|---|---|---|---|---|---|---|
| Parameter | Estimate | Lower | Upper | *t* | df | *p* | Std Est | Lower | Upper |
| (Intercept) | 0.38 | 0.08 | 0.69 | 2.54 | 111.28 | 0.013 | 0.11 | -0.06 | 0.28 |
| Language | -0.15 | -0.45 | 0.15 | -0.98 | 1325.00 | 0.329 | -0.08 | -0.25 | 0.08 |
| Vocoded | -0.01 | -0.31 | 0.29 | -0.08 | 1325.00 | 0.933 | -0.01 | -0.18 | 0.16 |
| Vocoded + Blurring | -0.73 | -1.03 | -0.43 | -4.76 | 1325.00 | $< 0.001$ | -0.41 | -0.58 | -0.24 |
| Language×Vocoded | -0.29 | -0.72 | 0.13 | -1.36 | 1325.00 | 0.175 | -0.16 | -0.40 | 0.07 |
| Language×Vocoded + Blurring | 1.06 | 0.63 | 1.49 | 4.87 | 1325.00 | $< 0.001$ | 0.59 | 0.35 | 0.83 |

| Random Effects | Variance | SD | ICC | Model Metrics | |
|---|---|---|---|---|---|
| Within-Group Variance | 2.69 | (1.64) | | AIC | 5334.2 |
| Between-Group Variance | | | | BIC | 5376.0 |
| Random Intercept (Participant) | 0.43 | (0.65) | 0.37 | $R^2$ (Conditional) | 0.16 |
| | | | | $R^2$ (Marginal) | 0.03 |
| N Groups | | | | ICC | 0.14 |
| Participant | 38 | | | RMSE | 1.62 |
| N Observations | 1368 | | | | |

##### 6.8 Word Onsets - LP - $-90$ ms

Table 9: Model formula: $Z \sim$ Language $\times$ SpectralDegradation + (1 | Participant). Standardized parameters were obtained by fitting the model on a standardized version of the dataset. 95% Confidence Intervals (CIs) and p-values were computed using a Wald t-distribution approximation.

| | | 95% CI | | | | | | 95% CI | |
|---|---|---|---|---|---|---|---|---|---|
| Parameter | Estimate | Lower | Upper | $t$ | df | $p$ | Std Est | Lower | Upper |
| (Intercept) | -0.42 | -0.66 | -0.19 | -3.53 | 92.61 | 0.001 | -0.05 | -0.22 | 0.13 |
| Language | 0.23 | 0.01 | 0.46 | 2.06 | 1325.00 | 0.040 | 0.17 | 0.01 | 0.33 |
| Vocoded | -0.60 | -0.83 | -0.38 | -5.34 | 1325.00 | $< 0.001$ | -0.44 | -0.61 | -0.28 |
| Vocoded + Blurring | 0.00 | -0.22 | 0.22 | -0.01 | 1325.00 | 0.994 | 0.00 | -0.16 | 0.16 |
| Language×Vocoded | 0.69 | 0.37 | 1.00 | 4.29 | 1325.00 | $< 0.001$ | 0.50 | 0.27 | 0.73 |
| Language×Vocoded + Blurring | 0.19 | -0.12 | 0.51 | 1.22 | 1325.00 | 0.224 | 0.14 | -0.09 | 0.37 |

| Random Effects | Variance | SD | ICC | Model Metrics | |
|---|---|---|---|---|---|
| Within-Group Variance | 1.46 | (1.21) | | AIC | 4508.8 |
| Between-Group Variance | | | | BIC | 4550.6 |
| Random Intercept (Participant) | 0.3 | (0.55) | 0.41 | $R^2$ (Conditional) | 0.22 |
| | | | | $R^2$ (Marginal) | 0.06 |
| N Groups | | | | ICC | 0.17 |
| Participant | 38 | | | RMSE | 1.19 |
| N Observations | 1368 | | | | |

##### 6.9 Word Onsets - LP - 420 ms

Table 10: Model formula: $Z \sim$ Language $\times$ SpectralDegradation + (1 | Participant). Standardized parameters were obtained by fitting the model on a standardized version of the dataset. 95% Confidence Intervals (CIs) and p-values were computed using a Wald t-distribution approximation.

| | | 95% CI | | | | | | 95% CI | |
|---|---|---|---|---|---|---|---|---|---|
| Parameter | Estimate | Lower | Upper | $t$ | df | $p$ | Std Est | Lower | Upper |
| (Intercept) | 1.43 | 1.17 | 1.68 | 11.10 | 94.88 | 0.00 | 0.70 | 0.54 | 0.86 |
| Language | -1.27 | -1.51 | -1.03 | -10.37 | 1325.00 | 0.00 | -0.81 | -0.97 | -0.66 |
| Vocoded | -0.48 | -0.72 | -0.24 | -3.89 | 1325.00 | 0.00 | -0.30 | -0.46 | -0.15 |
| Vocoded + Blurring | -1.52 | -1.76 | -1.28 | -12.43 | 1325.00 | 0.00 | -0.97 | -1.13 | -0.82 |
| Language×Vocoded | 0.24 | -0.10 | 0.58 | 1.37 | 1325.00 | 0.17 | 0.15 | -0.07 | 0.37 |
| Language×Vocoded + Blurring | 1.02 | 0.68 | 1.36 | 5.90 | 1325.00 | 0.00 | 0.65 | 0.44 | 0.87 |

| Random Effects | Variance | SD | ICC | Model Metrics | |
|---|---|---|---|---|---|
| Within-Group Variance | 1.71 | (1.31) | | AIC | 4721.8 |
| Between-Group Variance | | | | BIC | 4763.5 |
| Random Intercept (Participant) | 0.34 | (0.58) | 0.41 | $R^2$ (Conditional) | 0.30 |
| | | | | $R^2$ (Marginal) | 0.16 |
| N Groups | | | | ICC | 0.17 |
| Participant | 38 | | | RMSE | 1.29 |
| N Observations | 1368 | | | | |

### 7 Exploratory Analysis of Comprehension of Degraded Speech

#### 7.1 Neural Contrasts Across Experimental Conditions

| Reference | Comparison |
|---|---|
| Unprocessed English | Unprocessed Dutch |
| Vocoded English | Vocoded Dutch |
| Vocoded + Blurring English | Vocoded + Blurring Dutch |
| Unprocessed English | Vocoded English |
| Unprocessed English | Vocoded + Blurring English |
| Vocoded English | Vocoded + Blurring English |
| Unprocessed Dutch | Vocoded Dutch |
| Unprocessed Dutch | Vocoded + Blurring Dutch |
| Vocoded Dutch | Vocoded + Blurring Dutch |

Table 11: Pairwise contrasts between listening conditions for calculating neural contrasts in the exploratory analysis of comprehension of vocoded + blurring English.

##### 7.2 Neural Contrasts Correlating to Comprehension of Vocoded + Blurring English

| | Cluster | | | Time (ms) | | | | Rho | | |
|---|---|---|---|---|---|---|---|---|---|---|
| Feature | ROI | Reference | Comparison | Start | Max | End | Length | Sum | Max | $p_{cluster}$ |
| EnvDeriv | CF | Voc. Du. | Voc. Blur. Du. | 0 | 40 | 70 | 70 | 1.68 | 0.47 | < 0.0001 |
| EnvDeriv | LF | Voc. En. | Voc. Blur. En. | -50 | -10 | 20 | 70 | 1.67 | 0.48 | 0.0020 |
| EnvDeriv | RF | Voc. En. | Voc. Du. | 10 | 40 | 80 | 70 | 1.74 | 0.52 | < 0.0001 |
| EnvDeriv | RF | Voc. Blur. En. | Voc. Blur. Du. | 250 | 280 | 410 | 160 | 4.85 | 0.48 | < 0.0001 |
| EnvDeriv | CP | Unproc. En. | Voc. Blur. En. | 300 | 350 | 370 | 70 | 1.63 | 0.5 | 0.0020 |
| EnvDeriv | RP | Unproc. Du. | Voc. Du. | 20 | 70 | 120 | 100 | 2.94 | 0.47 | < 0.0001 |
| Envelope | CF | Unproc. En. | Unproc. Du. | 60 | 100 | 130 | 70 | 1.49 | 0.41 | 0.0010 |
| Envelope | CF | Voc. En. | Voc. Blur. En. | 100 | 120 | 180 | 80 | 1.87 | 0.42 | 0.0020 |
| Envelope | LF | Voc. Blur. En. | Voc. Blur. Du. | 350 | 380 | 410 | 60 | 1.17 | 0.44 | 0.0030 |
| Envelope | CP | Unproc. En. | Unproc. Du. | 210 | 240 | 280 | 70 | 1.46 | 0.41 | 0.0020 |
| Envelope | LP | Voc. Blur. En. | Voc. Blur. Du. | 220 | 260 | 300 | 80 | 1.85 | 0.41 | < 0.0001 |
| Envelope | RP | Unproc. En. | Unproc. Du. | 220 | 290 | 320 | 100 | 2.66 | 0.43 | < 0.0001 |
| Envelope | RP | Unproc. En. | Voc. En. | -10 | 30 | 60 | 70 | 1.66 | 0.47 | < 0.0001 |
| PhonemeOnsets | LF | Unproc. En. | Voc. En. | 80 | 160 | 290 | 210 | 6.32 | 0.43 | < 0.0001 |
| PhonemeOnsets | RF | Voc. En. | Voc. Blur. En. | 500 | 530 | 600 | 100 | 2.59 | 0.4 | < 0.0001 |
| PhonemeOnsets | LP | Unproc. En. | Voc. Blur. En. | 80 | 160 | 210 | 130 | 3.66 | 0.44 | < 0.0001 |
| PhonemeOnsets | RP | Voc. En. | Voc. Blur. En. | 510 | 580 | 600 | 90 | 2.42 | 0.45 | < 0.0001 |
| SpectralFlux | CP | Unproc. En. | Unproc. Du. | 380 | 420 | 440 | 60 | 1.16 | 0.49 | 0.0040 |
| SpectralFlux | CP | Voc. Du. | Voc. Blur. Du. | -50 | -10 | 40 | 90 | 2.73 | 0.57 | < 0.0001 |
| SpectralFlux | RP | Voc. Blur. En. | Voc. Blur. Du. | 50 | 90 | 120 | 70 | 1.65 | 0.48 | < 0.0001 |
| SpectralFlux | RP | Unproc. Du. | Voc. Blur. Du. | 60 | 90 | 120 | 60 | 1.09 | 0.41 | 0.0060 |
| SpectralVariation | RF | Unproc. Du. | Voc. Du. | 110 | 160 | 190 | 80 | 1.88 | 0.46 | 0.0010 |
| SpectralVariation | CP | Unproc. En. | Unproc. Du. | 170 | 210 | 240 | 70 | 1.44 | 0.42 | 0.0010 |
| SpectralVariation | LP | Voc. En. | Voc. Blur. En. | 240 | 280 | 320 | 80 | 2.04 | 0.5 | < 0.0001 |
| SpectralVariation | RP | Unproc. En. | Voc. En. | 150 | 190 | 240 | 90 | 2.24 | 0.45 | < 0.0001 |
| SyllableOnsets | CF | Voc. Blur. En. | Voc. Blur. Du. | 500 | 570 | 600 | 100 | 2.84 | 0.45 | < 0.0001 |
| WordOnsets | CF | Unproc. Du. | Voc. Du. | 350 | 440 | 500 | 150 | 4.74 | 0.44 | < 0.0001 |
| WordOnsets | CP | Unproc. En. | Unproc. Du. | 100 | 160 | 240 | 140 | 4.06 | 0.43 | < 0.0001 |
| WordOnsets | CP | Voc. Blur. En. | Voc. Blur. Du. | 490 | 560 | 600 | 110 | 3.06 | 0.42 | < 0.0001 |
| WordOnsets | CP | Voc. Du. | Voc. Blur. Du. | 400 | 450 | 560 | 160 | 4.75 | 0.47 | < 0.0001 |

Table 12: Neural contrasts surviving cluster-based permutation testing that individually predict comprehension of vocoded + blurring English. The significance of clusters is identified using mass permutation tests with 5000 iterations. Abbreviations: CF Central Frontal; LF Left Frontal; RF Right Frontal; CP Central Posterior; LP Left Posterior; RP Right Posterior. The threshold of significance for multiple comparisons over six regions of interest is set at $p_{cluster} \leq 0.008$.

#### 7.3 Neural Contrasts Contributing to Most Predictive Principal Component Analysis Solutions

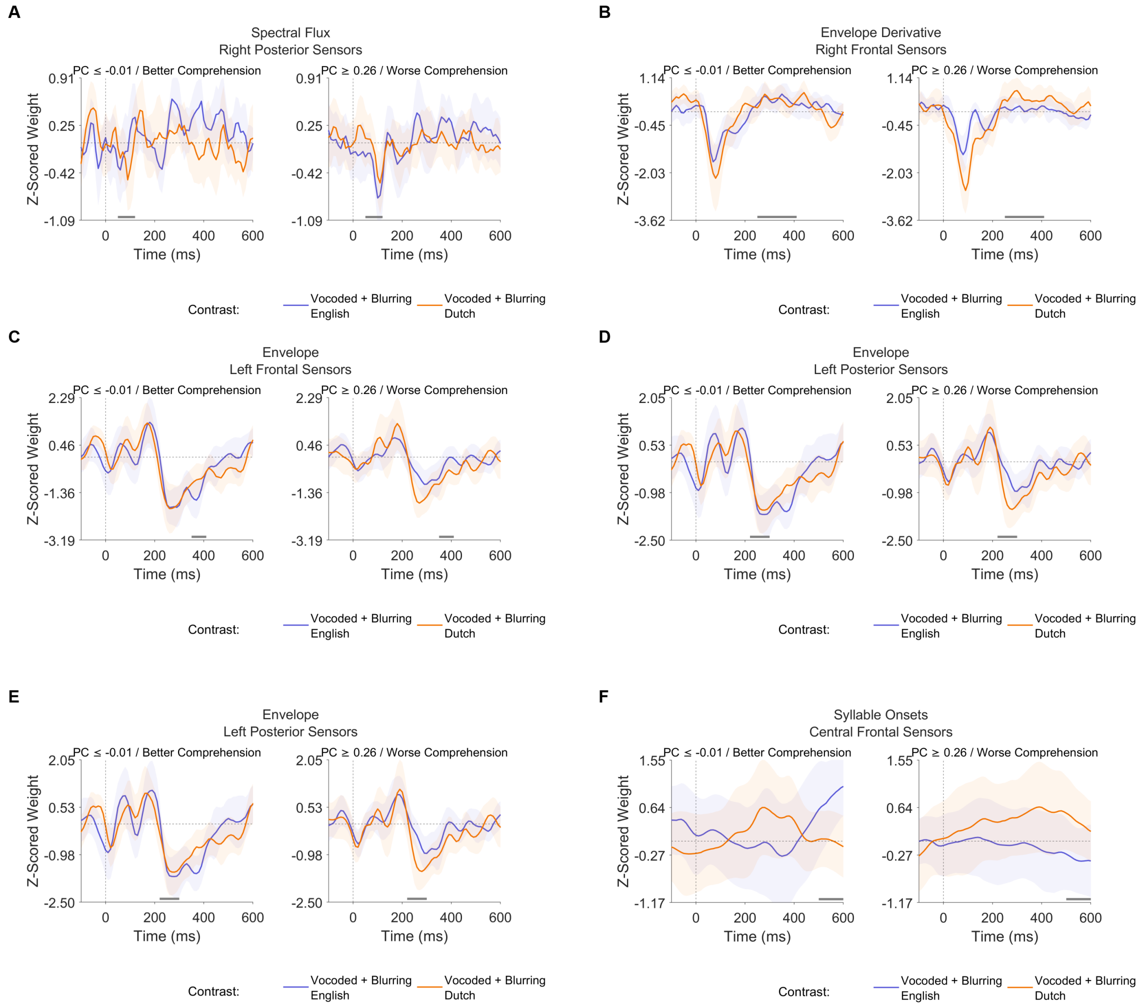


Figure 14: Encoding weights associated with the neural contrasts contributing most frequently to the top 5%-ranked PCA solutions, descriptively grouped by median-split principal component score. Grey bars indicate temporal regions where a correlation with comprehension of vocoded + blurring English for that neural contrast was found according to cluster-based permutation testing (5,000 iterations). Traces and shaded regions indicate the group mean (averaged over sensors) and 95% confidence intervals of the mean.

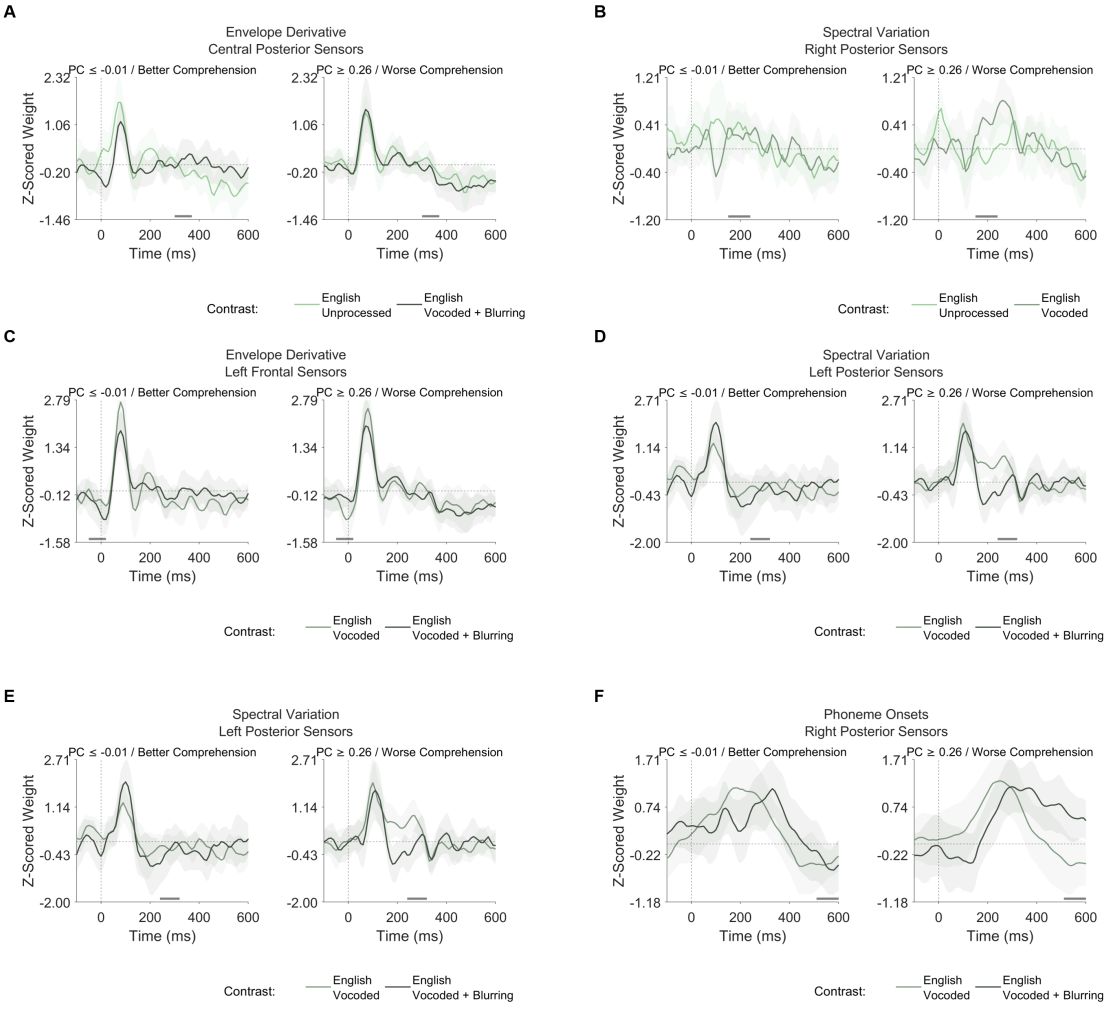


Figure 15: Encoding weights associated with the neural contrasts contributing most frequently to the top 5%-ranked PCA solutions, descriptively grouped by median-split principal component score. Grey bars indicate temporal regions where a correlation with comprehension of vocoded + blurring English for that neural contrast was found according to cluster-based permutation testing (5,000 iterations). Traces and shaded regions indicate the group mean (averaged over sensors) and 95% confidence intervals of the mean.

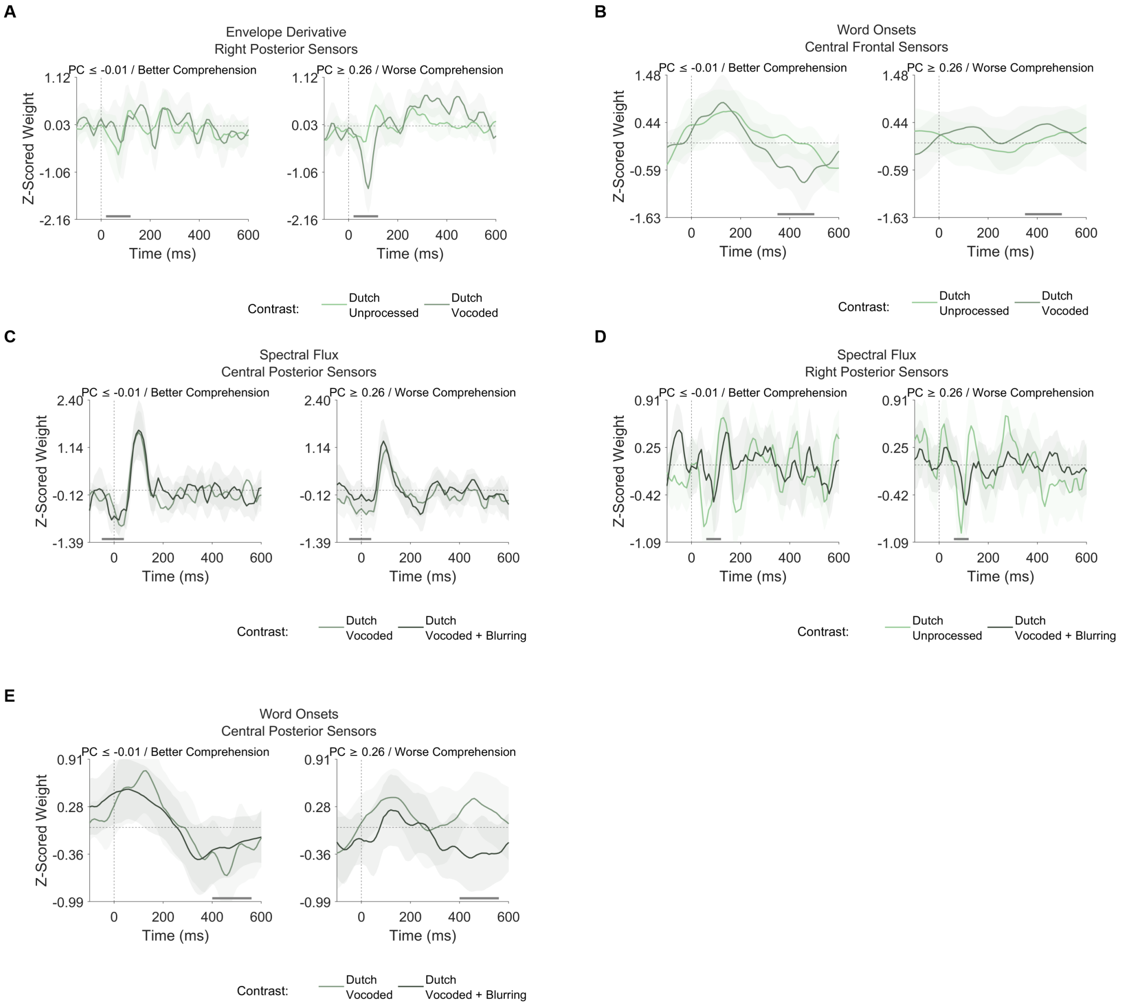


Figure 16: Encoding weights associated with the neural contrasts contributing most frequently to the top 5%-ranked PCA solutions, descriptively grouped by median-split principal component score. Grey bars indicate temporal regions where a correlation with comprehension of vocoded + blurring English for that neural contrast was found according to cluster-based permutation testing (5,000 iterations). Traces and shaded regions indicate the group mean (averaged over sensors) and 95% confidence intervals of the mean.

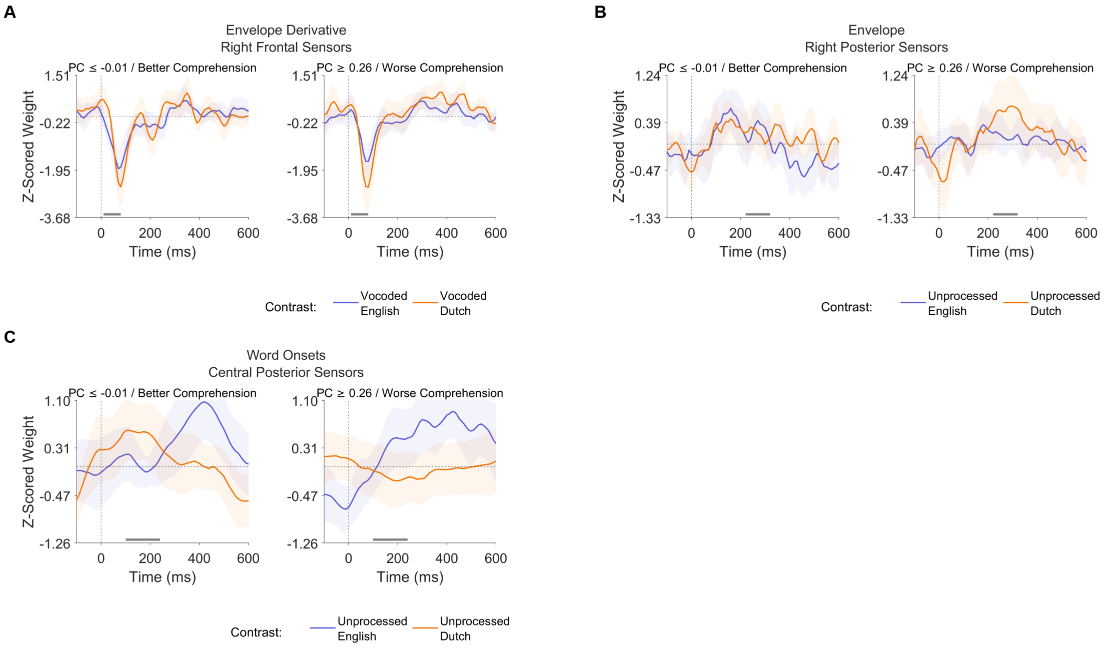


Figure 17: Encoding weights associated with the neural contrasts contributing most frequently to the top 5%-ranked PCA solutions, descriptively grouped by median-split principal component score. Grey bars indicate temporal regions where a correlation with comprehension of vocoded + blurring English for that neural contrast was found according to cluster-based permutation testing (5,000 iterations). Traces and shaded regions indicate the group mean (averaged over sensors) and 95% confidence intervals of the mean.
